# Exploring For Gloss: Active Exploration in Visual Material Perception

**DOI:** 10.1101/2024.07.09.602662

**Authors:** Lisa P.Y. Lin, Knut Drewing, Katja Doerschner

## Abstract

Image motion contributes to the perception of visual material properties, and motion signals are generated during active exploration. However, little is known about how specific perceptual tasks influence the actions that generate these cues. In an experiment using virtual reality and real-time hand tracking, we investigated how the demands of perceptual tasks (e.g., judging gloss or lightness) shape exploratory behaviours. Participants either observed or actively explored objects varying in gloss and lightness while performing a matching task. We analysed how their exploration patterns varied based on the tasks. Using the same stimuli in both tasks, we found that participants explored objects more extensively when judging gloss than when judging lightness. These findings suggest a strategic prioritisation of relevant cues for gloss judgments, with participants using larger movements and object rotation to enhance viewing perspectives and highlight detection. Our findings show that exploration behaviours are task-dependent, with actions adapted to the demands of the perceptual task at hand.

---

The perception of material properties is integral to our daily lives, as it helps us understand the qualities of the objects we encounter and guides how we interact with them. Every object and surface we see or touch is made of some material, each possessing distinct characteristics such as wetness, roughness, glossiness and softness. Being able to discern these material properties allows us to make practical decisions, such as determining whether an object is slippery, fragile, or durable, and to adapt our actions accordingly. For instance, choosing a metal railing for stability or adjusting our grip on a fragile object to avoid breakage (Buckingham, Cant & Goodale, 2009; Buckingham, Ranger & Goodale, 2011; Paulun, Gegenfurtner, Goodale & Fleming, 2016; Klein et al, 2021). Hence, the ability to perceive and recognise material properties of objects not only facilitates our immediate decisions but also shapes our actions.

Humans can readily recognise most objects’ material properties and almost instantaneously infer functional properties just by looking at them (Adelson, 2001; Fleming, 2014; Sharan, Rosenholtz & Adelson, 2009; Wiebel, Valsecchi & Gegenfurtner, 2013). Prior studies on visual perception of materials have examined and identified several physical factors and visual cues that influence our perception of optical material properties, such as gloss, translucency, and lightness in static scenes (for a review, see Anderson, 2011; Gigilashvili, Thomas & Hardeberg, 2021; Chadwick & Kentridge, 2015; Murray, 2021). For instance, research on gloss perception has found that object shape and surface geometry (Vangorp, Laurojssen & Dutré, 2007; Ho, Landy & Maloney, 2008; Marlow, Kim & Anderson, 2012; Marlow & Anderson, 2013), illumination conditions (Fleming, Dror & Adelson, 2003; Doerschner, Boyaci & Maloney, 2010 a,b; Motoyoshi & Mataba, 2012; Olkkonen & Brainard, 2010), and specular highlights properties like their shape, intensity, location and the orientation between highlights and shading have an impact on perceived gloss (Berzhanskaya, Swaminathan, Beck & Mingolla, 2005; Beck & Prazdny, 1981; Kim, Marlow & Anderson, 2011). In dynamic scenes, researchers have also demonstrated the importance of motion cues in perceiving optical material qualities (e.g. Doerschner et al., 2011; Wendt, Faul, Ekroll & Mausfeld, 2010; Sakano & Ando, 2010; Lichtenauer, Schuetz & Zolliker, 2013). Similarly, lightness perception has been found to be influenced by factors such as direction and intensity of illumination (Kobayashi & Morikawa, 2019; Gilchrist et al., 1999; Toscani, Zdravković & Gegenfurtner, 2016), contextual information like contrast with surrounding elements and background (Hess & Pretori, 1970; Allred & Brainard, 2013), as well as surface reflectance, orientation, texture, and shape (Toscani, Valsecchi & Gegenfurtner, 2017; Schmid & Anderson, 2014; Boyaci, Maloney & Hersh, 2003; Knill & Kersten, 1991; Pessoa, Mingolla & Arend, 1996). These examples highlight the visual system’s reliance on different *visual cues*, yielded by intrinsic factors like surface geometry and object size, and extrinsic factors, such as illumination properties and viewpoints, to infer and estimate visual material properties.

Yet, humans do not passively rely on these visual cues. Instead, they actively seek information that is relevant to the task at hand (Rothkopf, Ballard & Hayhoe, 2007). Research on eye movement patterns have shown that people strategically direct their gaze toward areas of an object that offer the most relevant information for judging its visual material qualities. For example, when assessing the lightness of matte objects, observers tend to focus on the brightest part of the surface and base their judgments on those regions (Toscani, Valsecchi & Gegenfurtner, 2013; 2017). If observers are forced to focus on a specific area, the lightness of that area will influence their lightness judgments of the entire surface. Interestingly, the selection of the fixation region is also influenced by surface properties: when judging the lightness of glossy surfaces, observers prioritise regions adjacent to the highlights over the brightest parts, as highlights are not diagnostic of surface lightness (Toscani et al., 2013). In another eye-tracking study, Toscani, Yücel and Doerschner (2019) found that participants focused more on gloss-relevant image motion cues (Doerschner et al., 2011) during gloss judgments than during speed judgments.

Taken together, the research on visual cues suggests that visual sampling strategies are optimised for specific tasks and are modulated by materials properties, with observers looking at areas in static and dynamic scenes that contain the most task-relevant information. However, in real-world settings we judge the qualities of materials not only by looking at objects-but also by manually exploring and interacting with them (Figure 1).

**Figure 1.**
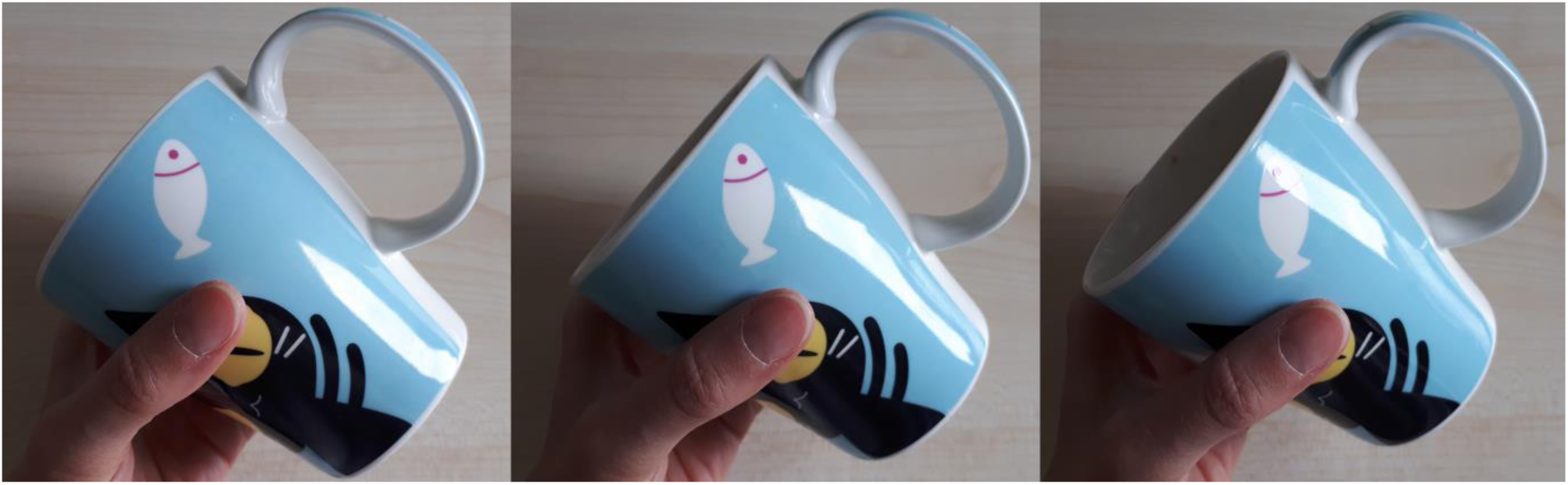
Active exploration of objects and materials. Active exploration of objects and materials produces visual cues that aid subsequent visual recognition and estimation tasks. For instance, when observing glossy object, people often turn them in their hands to see how the highlights move across the surface. This suggest that the movement of these visual features during motion or changes in perspectives may convey useful information.

This interaction potentially provides us with the cues that the visual system learns to use for estimating object properties (e.g. Adolph, Eppler & Gibson, 1993, Metta & Fitzpatrick, 2003, Smith, Jayaraman, Clerkin & Yu, 2015). This is not only true for mechanical material properties but also for the perception of optical material properties, where interaction is adapted to extract the most useful information based on the perceptual goal, as illustrated in Figure 1. For example, Gigilashvili and colleagues (2018; 2019; 2021) observed that when participants were asked to assess an object’s translucency or glossiness, they adapted their actions to the task, where they manipulate the object or change their viewpoint to gather the most relevant visual information. This suggests that exploratory actions, such as object manipulation and head movement, are not merely incidental, but are driven by the perceptual demands of the task.

Previous work points to the idea, that motion signals, generated through such interactions, indeed play an important role in determining material appearance (e.g. specular flow or highlight motion for glossiness, see e.g. Doerschner et al., 2011; Wendt et al., 2010; Sakano & Ando, 2010). For example, Sakano and Ando (2010) investigated the role of motion parallax on gloss perception. They found that dynamic cues, particularly the relations hip between the observer’s movement and the resulting motion of specular reflections, significantly enhanced the perception of glossiness. Their results showed that it is not simply the act of moving one’s head that aids the perception of gloss, but rather the temporal changes in the retinal image caused by motion. In their experiment, participants observed static stimuli (where the visual information remained constant despite head movement) and dynamic stimuli (where the object changes according to head movement). The results showed that head movement alone, without corresponding changes in visual information, did not enhance gloss perception. This supports the idea the perceptual tasks, such as judging gloss, could benefit from motion-related information, - since mere multiframe observation (such as static images) may be insufficient to provide the same level of information. This raises the question, to what extent human exploratory hand movements are tuned to the visual perceptual task when estimating optical material properties. Interacting with an object through motion, whether by hand or head, seems to provide crucial information that the visual system needs to estimate properties such as gloss.

To answer this, one has to consider the potential information to be gained through interaction. For example, if the task is to judge the glossiness of a smooth bumpy object, much is to be gained by rotating the object with respect to the light source, because that would generate the type of highlight motion that has been shown to aid in assessing the glossiness of a surface. As illustrated in Figure 2A, rotating a glossy object around the vertical axis a small amount, e.g. 10 degrees. Measuring the change in pixel intensity between two frames, where the object has an angular difference of 10 degrees, (Figure 2C) we see that quite a few pixels change their intensity (brighter colours, denote larger pixel differences). Suppose now that the task is to judge the lightness of a similar, but matte object, as the one shown in Figure 2B. A rotation by the same amount (10 degrees) around the vertical axis would lead to much less change in pixel intensity (Figure 2D). Expanding the analysis to a larger rotation angle of 45 degrees (Supplementary Figure S1) confirms this pattern. For the glossy textured object (Figure S1A), the larger rotation causes even more significant pixel intensity changes, as shifting highlights become more pronounced (Figure S1D), increasing the information gained through movement. Conversely for the matte textured and non-textured objects (Figure S1B and C), even at 45 degrees, there is still much less change in pixel intensity (Figure S1E and F), again indicating less information is obtained from the movement. This relationship holds true not only for smooth surfaces but also for textured, bumpy objects. Therefore, the difference in visual information between glossy and matte surfaces is directly related to the presence of gloss. While gloss introduces dynamic, high-frequency changes as the object rotates, matte surfaces—whether smooth or finely textured—exhibit fewer changes, and the same pattern holds true regardless of the rotation angle. Interaction, particularly through rotation, is therefore essential for extracting material properties from glossy surfaces, while matte surfaces provide much less information through similar movement.

**Figure 2.**
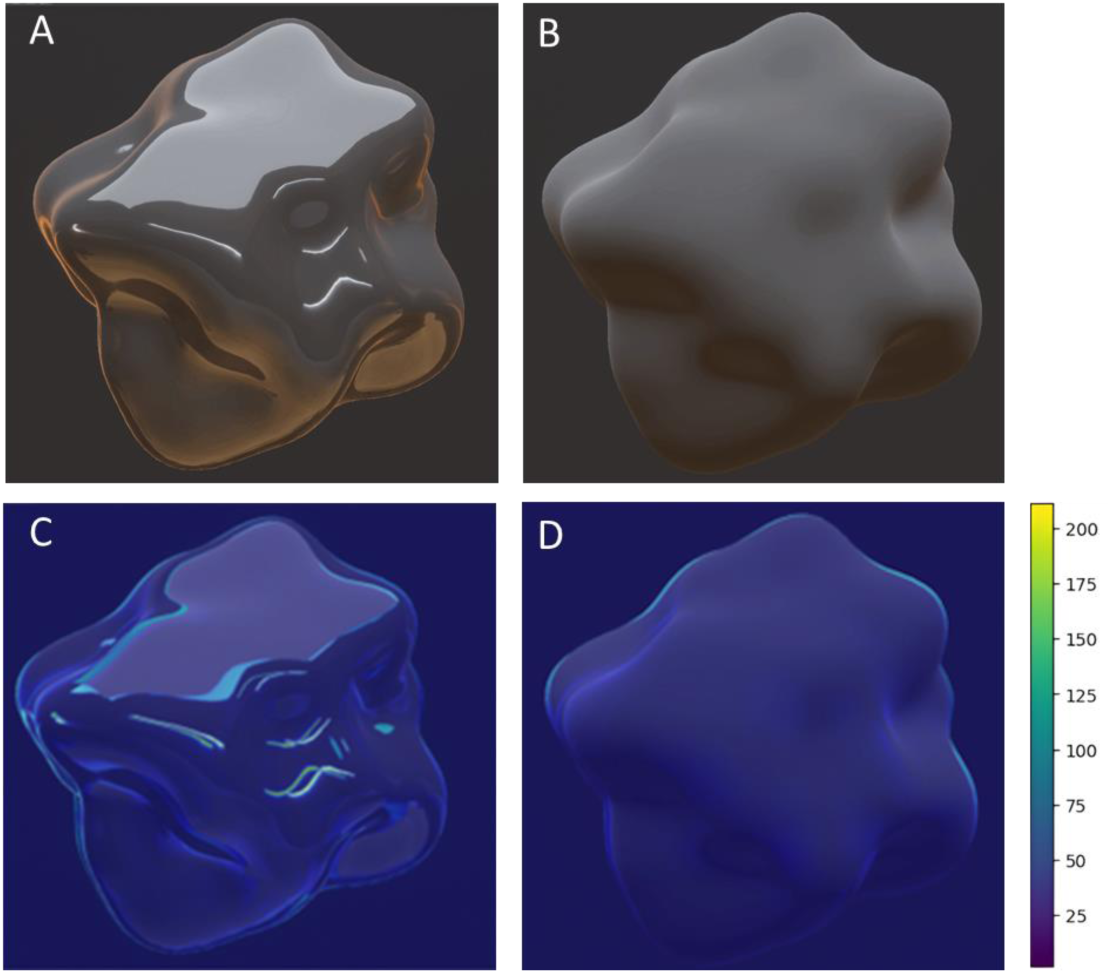
Absolute pixel intensity difference across frames. Example of absolute pixel intensity difference across frames. Panel A shows a rotating high-gloss object, and Panel B shows a rotating matte object. Panels C and D display the average absolute pixel intensity difference between two consecutive frames, where the objects differ by a 10-degree rotation. See text for details.

Would we find differential exploration behaviour as a function of perceptual task and material in an experiment? To test this general idea, we conducted a pilot experiment (see Supplementary materials, section A). Using different stimuli for gloss and lightness tasks, we found preliminary evidence for task-dependent exploration differences. Based on this pilot experiment, we optimised the design and developed an active exploration paradigm in virtual reality coupled with real-time hand tracking to investigate whether observers explore objects differently when judging gloss or lightness. Our approach paralleled Ho, Landy and Maloney (2008), who used conjoint measurement to examine gloss and surface texture perception. Similarly, we used stimuli that varied in both gloss and lightness, allowing us to assess task-driven exploration strategies in an interactive setting.

Participants were tasked with matching the gloss or lightness of an object to a set of seven comparison objects, and they either interacted with the object or observed the object while forming their judgments. We hypothesised that active interactions would provide useful information for judging visual gloss, leading to more rotational movements during gloss judgements than lightness judgements (Doerschner et al., 2011; Toscani et al., 2019). In contrast, the role of image motion in lightness judgments is less clear, as object motion can sometimes compromise lightness constancy (Wendt et al., 2010). We expected, when judging lightness, participants might selectively focus on certain areas, e.g. the brightest part of the object, which provides the most informative cues for their judgments (Toscani et al., 2013; 2017). Since the brightest region of a matte object will not change substantially when moving the object in hand, we expected fewer overall movements during lightness judgements.

## Methods

### Participants

Twenty-five participants (7 males) between 20 to 39 years of age, M_age_ = 28.40 years, SD_age_ = 5.08 years) were recruited from Giessen University. All had normal to corrected-to-normal vision and had no history of motor impairments. Participants provided informed consent and received compensation of (8€/h) for their participation. This study was approved by the ethics committee at Giessen University (LEK FB06) and conducted in accordance with the declaration of Helsinki (2013) except for preregistration. One participant was excluded due to technical issues during data collection, and another for not understanding the instructions.

### Stimuli

The stimuli consisted of 3D virtual objects rendered with varying levels of smoothness and albedo. We used perceptually distinct levels of smoothness and lightness. Levels were determined in a separate, additional pilot experiment (N = 5).

#### Shapes

Two novel 3D shapes (‘Glaven objects’; Phillips et al., 2009; 2016) were used as the target and comparison objects (Figure 3). The target object had an apparent diameter of 20 cm. When held at arm’s length, participants viewed the object at a visual angle of approximately 8–10 degrees. During the interactive condition, as participants moved the object closer or farther from themselves, this visual angle dynamically adjusted in response to the object’s distance.

**Figure 3.**
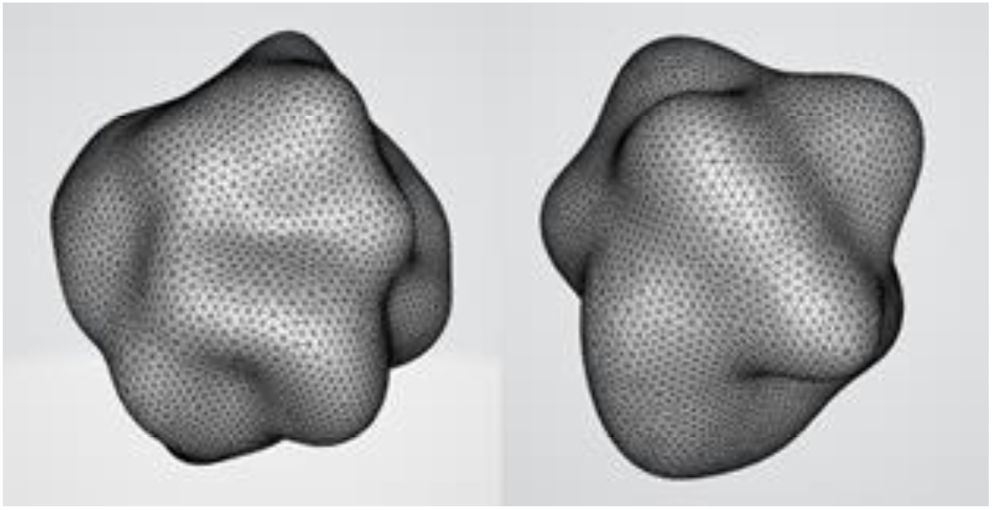
3D shapes used in the experiment. Left: target shape, right: comparison shape.

### Material

To generate the material for our stimulus sets, we utilised the Unity HDRP built-in Lit shader, changing the albedo and smoothness parameters. In Unity, the Albedo property determines a material’s base colour (RGB), and the smoothness scale controls the degree of blur in specular reflections. The RGB channel values range from 0 to 1, with greyscale colour resulting when the R(red), G (green), and B(blue) channel values are equal. For instance, black is represented as [0 0 0] and white as [1 1 1]. The smoothness scale operates from 0 to 1, where 0 represents a completely matte surface, and 1 signifies a flawlessly smooth surface that reflects light like a mirror.

The stimuli were created by combining four gloss levels and four Perlin noise-based lightness levels, to generate 16 unique stimuli representing all possible combinations (see Figure 4). For the gloss levels of the target objects, we used four smoothness values: 0.35, 0.55, 0.70, and 0.85, ranging from least glossy to most glossy. For the lightness levels of the target objects, we generated Perlin noise textures with adjusted contrast and brightness to create four different lightness levels.

**Figure 4.**
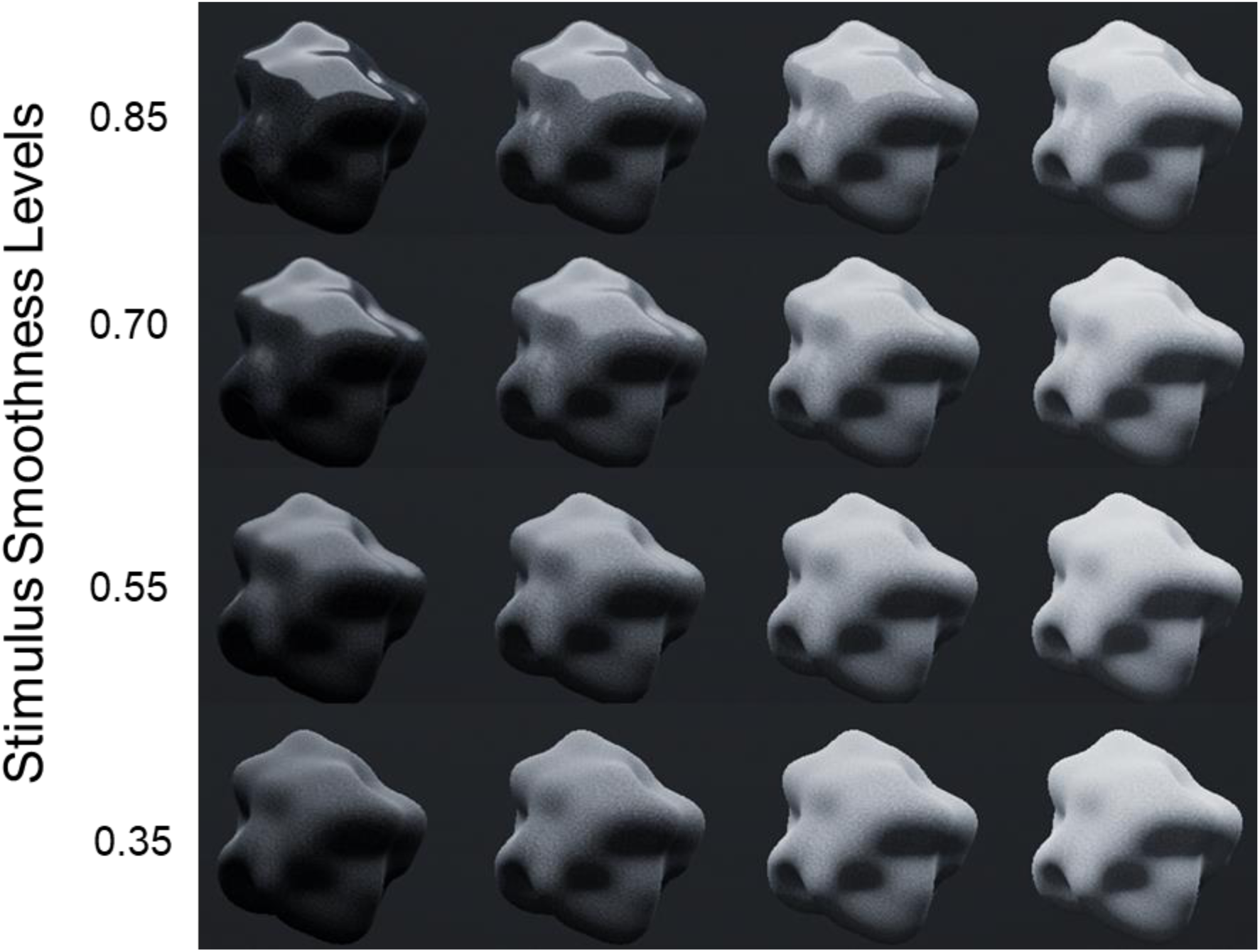
The 16 target objects used in all conditions. Objects are arranged from left to right in order of increasing lightness (darkest to lightest) and from bottom to top in order of increasing gloss (least glossy to most glossy). The estimated diffuse reflectance values for the four lightness levels, from lightest to darkest, were 58%, 47.5%, 32.5%, and 17.5%, respectively. Specular reflectance, based on smoothness levels for the four gloss levels, was approximated as 35%, 55%, 70%, and 85%, from least to most glossy,

The decision to use Perlin noise-based textures for the lightness levels arose from the need to use the same stimuli for both gloss and lightness judgments. Since darker surfaces often appear glossier, it was essential to include a range of lightness values. Perlin noise textures allowed us to represent these variations, ensuring the stimuli were suitable for both gloss and lightness judgements.

#### Perlin Noise Texture Generation

The Perlin noise textures used in the experiment are generated using the *Mathf.PerlinNoise* function in Unity, which produces values between 0 and 1. These values are scaled by a multiplier of 0.25 to reduce the contrast, and a bias is added to control the overall brightness. The pixel values for each texture are calculated using the formula:

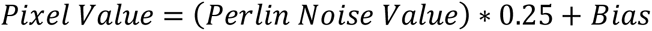

By adjusting the bias, we create different mean levels of lightness. The four bias values used to generate the four lightness levels were: 0.455 for lightness level 1 (lightest), 0.35 for lightness level 2, 0.2 for lightness level 3, and 0.05 for lightness level 4 (darkest), see Figure 5.

**Figure 5.**
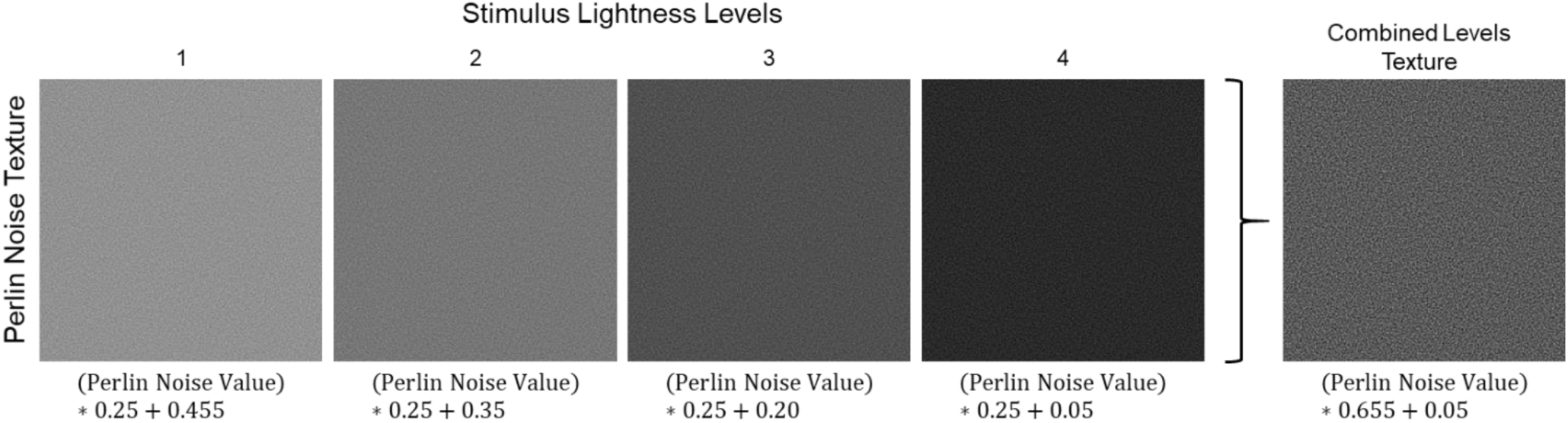
Perlin noise textures generated for the different lightness levels. From left to right, the textures are arranged in decreasing lightness (lightest to darkest). The texture on the far right is the combined levels texture, which encompasses the full range of lightness levels used in the target objects and is used for the gloss comparison objects.

#### Combined Levels texture

Additionally, we generated a texture that encompasses the full brightness range of all lightness levels for the gloss comparison objects. This texture was created using a multiplier of 0.655 and a bias of 0.05, covering the brightness range from 0.05 to 0.705 (see Figure 5 far right panel).

#### Comparison objects

For the gloss comparison objects (see Figure 6 top panel), we used the combined levels texture and varied the smoothness values (0.05, 0.25, 0.50, 0.65, 0.75, 0.85, 0.95) to create seven different gloss levels, ranging from least glossy to most glossy. For the lightness comparison objects (see Figure 6 bottom panel), we set the smoothness value to 0 (completely matte) and used RGB values of 0.60, 0.50, 0.40, 0.30, 0.20, 0.09, and 0.01 to generate seven lightness levels, organised from lightest to darkest.

**Figure 6.**
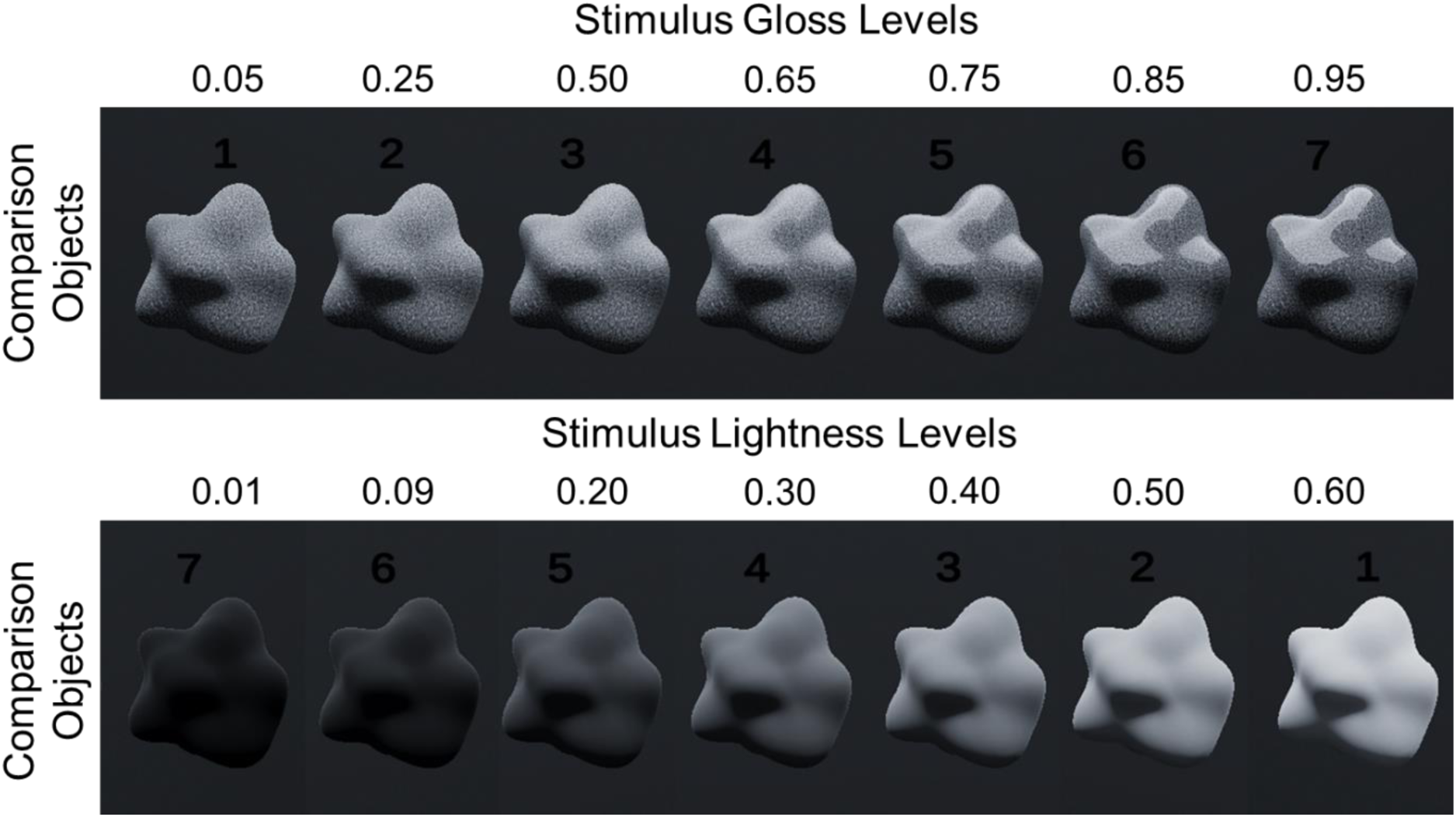
Comparison objects used in this experiment. Top panel: Gloss condition comparison objects. The diffuse reflectance was approximated by calculating the average brightness (representing the albedo of the texture), resulting in an estimated 37.75%. Specular reflectance was based on smoothness levels across seven gloss levels (from least to most glossy): 5%, 25%, 50%, 65%, 75%, 85%, and 95%. Bottom panel: Lightness condition comparison objects. The estimated diffuse reflectance values, from lightest to darkest, were 60%, 50%, 40%, 30%, 20%, 9%, and 1%. Smoothness was set to 0 in Unity’s smoothness scale, making the objects fully matte with negligible specular reflectance.

#### Luminance measurements

We measured the approximate physical luminance and chromaticity values for each material using a spectroradiometer (KONIKA MINOLTA CS-2000). Detailed values can be found in Supplementary Tables S1 and 2. Measurements were performed using the same 3D object used as the target object in the experiment, similar to how they appeared to participants in the virtual environment. Additionally, we measured the materials on a flat cube.

#### Light field and environmental lighting

The default HDRI sky was used for the virtual environment to provide ambient lighting based on high-dynamic-range images (HDRI). The Default HDRI Sky texture was imported into Unity as a “Default” texture type with the texture shape set to “Cube.” Mapping was set to “Auto”, and Convolution type was set to “None”. Wrap mode was set to “Repeated,” with filter mode as Bilinear. Default settings included a maximum texture size of 2048 pixels, resize algorithm set to “Mitchell” for high-quality resizing, Format set to “Automatic,” and Compression at “Normal Quality”. In addition to the default HDRI sky texture, we used two area lights: one area light (3.86m x 2.58m, 600 lumens, colour = white [RGB 1, 1, 1]) positioned 1.836m above the virtual table at an angle of 90 degree relative to observer’s view point, and a second area light (same size and colour, 45 lumens) placed 1.103m from the table, facing the virtual objects (same orientation as observer’s viewpoint).

### Apparatus & software

The experiment programme and environment were created by us using the Unity 3D Gaming Engine (version 2021.3.9f1) High-Definition Render Pipeline (HDRP) with the Ultraleap plugin. Participants wore a VIVE Pro Eyes head-mounted display (HMD) that displayed a stereoscopic image of the virtual environment with a resolution of 1440 x 1600 pixels per eye (2880 x 1600 pixels combined), with a field of view of 110 degrees and a frame rate of 90Hz. The participant’s head movements were tracked, updating their perspective as they explored the virtual environment by turning their head. The position of participants’ arms and hands was tracked using an Ultraleap Leap motion controller mounted on the front of the HMD using a custom 3D-printed mount. The Leap Motion controller consisted of two near-infrared cameras that track the user’s hand movement with a resolution of 640 x 240 pixels per camera and a framerate of 60 frames per second. The movement and position of the participant’s tracked hands were mapped onto the virtual hand in real-time so that the movement of the virtual hand was congruent with the movement of the participant’s actual hand. The avatar hands utilised were sourced from the Ultraleap Gemini (V5.2) SDK’s hands module (see Figure 7).

**Figure 7.**
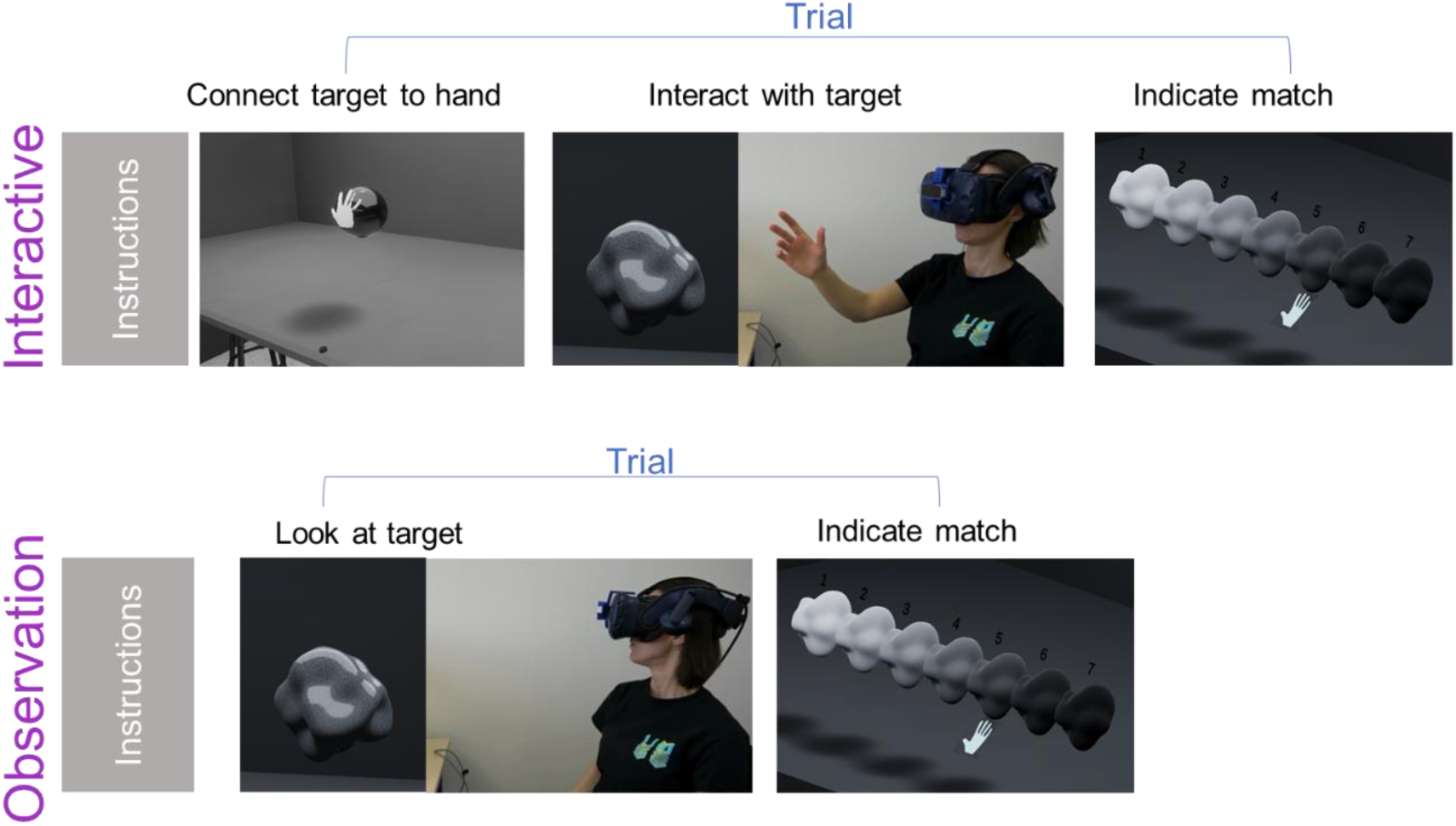
Trial sequences for interactive and observation conditions. Interactive: A chrome sphere indicates the starting hand position before the onset of exploration, and the virtual hand participants used to interact with the virtual object in the interactive condition. The participant could freely explore the object that was now yoked to the participant’s hand. Once exploration was completed, comparison objects would appear, and the participant indicated their match verbally. The response would be logged in by the experimenter, after which the next trial would start. Observation conditions: Same as the interactive conditions, except for the object-hand yoking procedure.

#### Scene Layout

The virtual environment featured a room with a grey wall and table where the experiment took place. A 3D camera, positioned at eye level to match the height of the target object, aligned with a black dot (2cm in diameter) placed on the table, which served as a reference point, helping participants establish a consistent egocentric position throughout the experiment.

In all conditions, the target objects were presented at the same location across trials, approximately 35.5cm from the reference point. However, the orientations of these objects were randomised to prevent participants from fixating on specific object regions during exploration or observation. The set of comparison objects were stationary and arranged in a straight row, centred at 35.5 cm from the reference point.

### Design and Procedure

In a single session, participants engaged in four experimental conditions: Interactive Gloss, Interactive Lightness, Observation Gloss, and Observation Lightness, the order of which was counterbalanced across participants. Each condition consisted of 16 trials, where each of the 16 stimuli was presented once in randomised order. The initial perspective of the target object was varied across ten possible orientations involving rotations along either the x or y-axis by 45°, 90°, 135°, 180° or 225°. The session’s duration averaged around 1 to 1.5 hours.

After providing their informed consent, participants were seated at a table and received instructions (Figure 7) for the experiment. They were encouraged to ask questions if anything was unclear. Participants were told that their task was to make matching judgments about either gloss or lightness: they would be presented with a target object, which they would interact with or observe for as long as they needed to, and then select the comparison object that best matches that target object in terms of the specified visual material quality (gloss or lightness). Once participants understood the task, they put on the VR headset, and the experiment began. Before each condition started, instructions were again displayed in the virtual environment.

In each trial, participants were first presented with a target object, depending on the condition (interactive or observation), participants either explored or observed the target object at their own pace. When ready to make their judgment, they notified the experimenter, which stopped the timer tracking their exploration or looking time. At that point, seven comparison objects appeared, each varying in gloss or lightness. Participants then select the comparison object that best matched the target object in terms of the specified material (i.e., gloss or lightness). The comparison objects were numbered, and participants indicated their selection by verbally stating the corresponding number to the experimenter. After the experimenter had recorded their responses, the comparison objects disappeared, and the experimenter pressed the enter key to start the next trial. The general procedure remained consistent for both material judgments (gloss/lightness). The procedure for the exploration conditions (interactive/observation) differed slightly and is outlined next:

#### Observation conditions

The procedure of this exploration condition was as outlined above, and participants were instructed to minimise body and head movements throughout this exploration condition.

#### Interactive conditions

In this exploration condition, participants were provided with a virtual hand, which was only visible when they were positioning their hand on the starting position at the beginning of each trial. Participants first saw a chrome sphere in front of them, and were instructed to place their virtual right hand on the sphere as if preparing to grasp it (Figure 7). This step ensured a uniform hand positioning before exploration began. Once their hand was properly positioned, a target object appeared and simultaneously, the virtual hand became invisible. The target object became attached to the palm of the now invisible virtual hand. Participants could move and rotate the target object by moving and rotating their physical hand. They could freely interact with the object as if they were manipulating it with their physical hand, with both rotational and translational movements possible. Additionally, their hand movements during explorations were captured by the Ultraleap Leap motion controller. This design ensured participants focused on the target object rather than their virtual hand, and prevented the hand from intersecting with the object mesh. After participants made their matching judgement, the comparison objects disappeared once the experimenter recorded their response. The chrome sphere and their virtual hand then reappeared, signalling the start of the next trial.

### Measurements

We recorded several hand movement parameters to assess potential differences in exploration between the gloss and lightness judgements conditions. We also assessed looking time during the observation condition.

#### Looking/Exploration time

We measured the time duration during which participants observed or explored the target stimulus before making their judgments, as exploration, whether visual or manual, is often driven by the need to extract and process task relevant information (e.g., Kaim & Drewing, 2011; Toscani et al., 2017; Philips et al., 2010). For the observation conditions, we measured the looking time, which refers to the duration (in seconds) between the onset of the target stimulus and the onset of the comparison objects. Whereas in the interactive conditions, we measured their exploration time, which is the time elapsed (in seconds) between the onset of the target stimulus (i.e., the start of exploration) and the onset of the comparison objects (i.e., the end of exploration). These measurements allow us to assess how long participants observed/explored the stimulus before forming their judgements.

#### Palm Distance

We measured the extent of hand movement made by participants during active exploration. To do this, we used the ‘palm position’ attribute recorded by the Leap Motion controller, which contains the 3-dimensional coordinates of the palm centre point (x, y, z, see Figure 8). We calculated the distance travelled by the palm during exploration by quantifying the distance between each set of coordinates in successive frames (*i,i+1*) using the following formula: 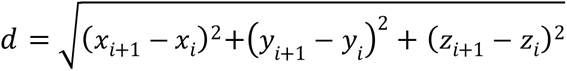. Following this, the palm distance value is computed as the sum of distances between all consecutive frames within a trial. This measurement enables us to evaluate the extent of translational movements made by participants during exploration.

**Figure 8.**
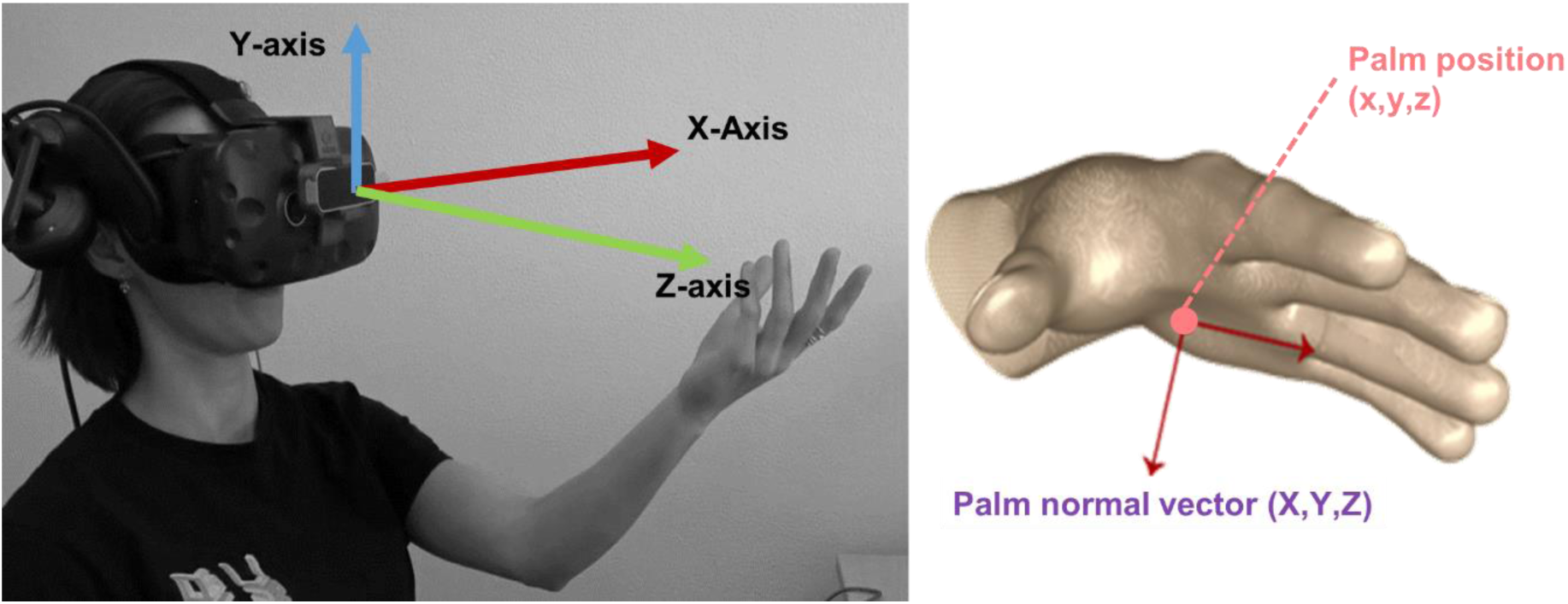
Leap motion coordinate system and hand movement parameters. Depiction of the Leap Motion coordinate system, illustrating the palm centre position and palm normal vector.

#### Palm Rotation

This measurement is determined using the ‘palm normal’ attribute captured by the Leap Motion controller. This attribute comprises the 3-dimensional coordinates of a unit vector (X,Y,Z) perpendicular to the plane formed by the palm of the hand. For instance, if the palm is flat and facing upwards, the vector points upwards (Figure 8). We use this to determine the rotation of the palm as it rotates during interaction with the stimulus. Similar to the cumulative palm movement, we first quantified the distance between each set of coordinates in successive frames using the formula: 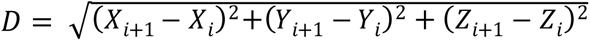. Then, we sum these distances across all consecutive frames, and finally convert this sum to cumulative rotational angle by multiplying it by*180*/π. Using the measurement, we can gauge the extent to which participants rotate their hand during exploration.

### Data analysis

#### Perceptual Judgments

Before analysing exploration behaviour, we first assessed whether participants performed the perceptual tasks reasonably. We calculated the mean perceived gloss and lightness for each gloss/lightness level across the two conditions (interactive and observation). An ANOVA was then conducted for each condition to determine whether participants could reliably differentiate between the stimulus levels. The results revealed significant main effects of stimulus levels (*ps* = <.001, pairwise comparisons: *p* =.008 to <.001), indicating that participants were performing the tasks as expected. Next, we conducted paired-sample t-tests to compare perceived gloss and lightness between the two exploration conditions (interactive vs. observation) to evaluate whether the condition influenced participants’ judgements.

#### Exploration behaviours/ parameters

The data (averaged for each participant for each perceptual task, gloss and lightness level) for each exploration parameter (exploration time, looking time, palm distance and palm rotation) were analysed separately with repeated measure ANOVAs. Bonferroni post-hoc pairwise comparisons were used to identify specific differences between main effects levels. Greenhouse-Geisser correction was applied whenever the sphericity assumption is violated.

#### Interobserver reliability

To evaluate the consistency of perceptual judgments made by participants in each condition, we also calculated Pearson’s correlation coefficients between all possible pairs of participants across all stimuli. Similarly, for exploration data, we computed Pearson’s correlation coefficients between all possible pairs of participants across all stimuli to assess their interobserver agreement within each task and exploration parameter. These parameters include exploration time, palm distance, and palm rotation, for interactive conditions and looking time for observation conditions.

## Results

### Exploration Behaviours

The main goal of the current experiment was to determine whether exploration behaviour differed between perceptual tasks (judging gloss vs. lightness) using the same set of stimuli. We assessed variations in exploration parameters across perceptual tasks, Figure 9 shows an overview of the results. Overall, we found that in the interactive conditions, participants spent more time and engaged in more extensive exploration during the gloss judgement task compared to the lightness judgement task. However, in the observation conditions, looking time did not differ between the two tasks.

**Figure 9.**
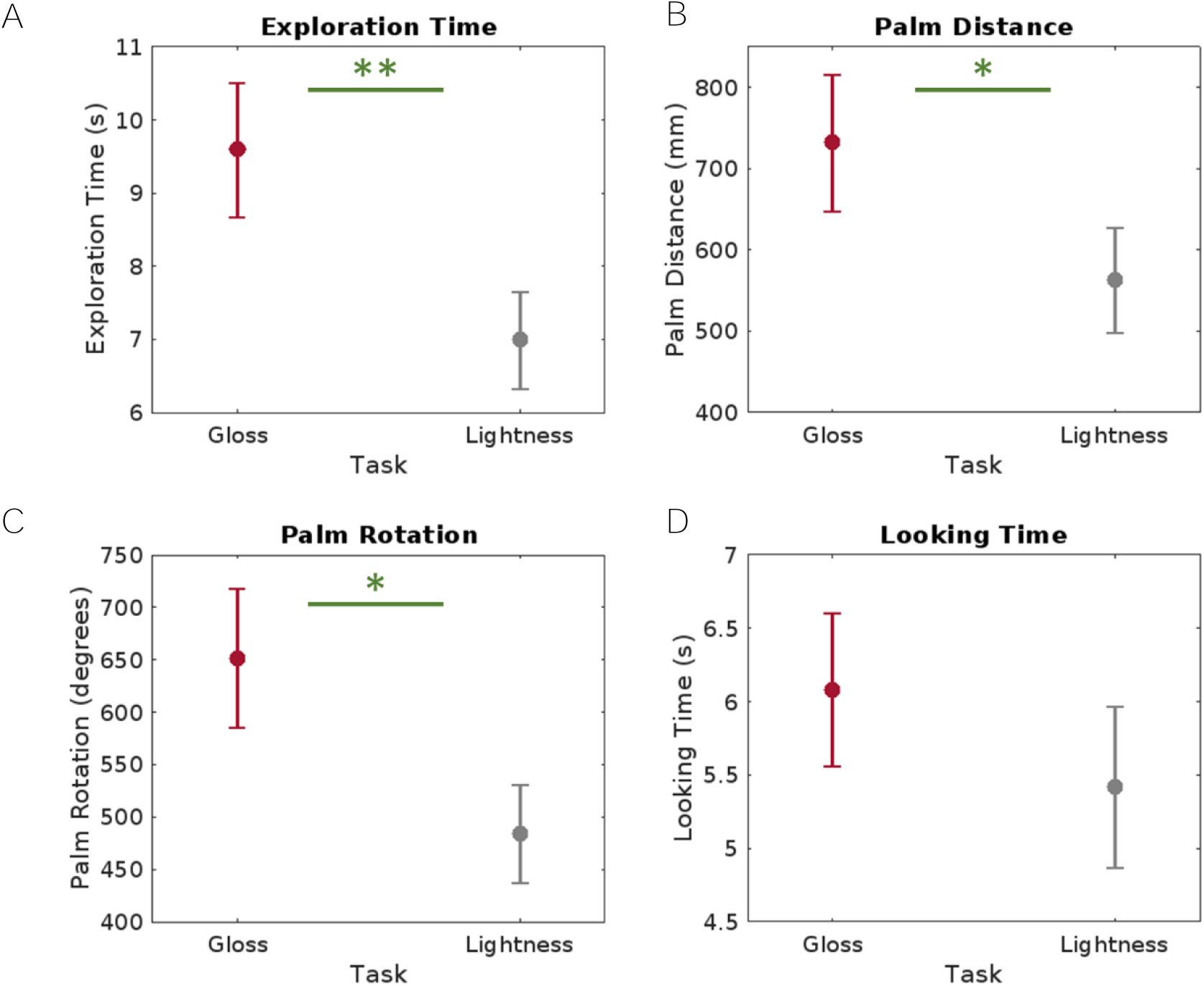
Results exploration parameters. A) Mean exploration time, B) Mean palm distance, C) Mean palm rotation, D) Mean looking time plotted as a function of perceptual task. Error bars represent 1 ± SE.

### Interactive conditions parameters

#### Exploration time

Overall, we found that participants spent longer time actively exploring the stimulus in the gloss condition (*M_gloss_* = 9.599, *SE* = .916) than in the light ness condition (*M_lightness_* = 6.991, *SE* = .662, *p* = .007). A repeated-measures 3-way ANOVA revealed a significant main effect of the perceptual task, *F*(1,22) = 8.975, *p* = .007, *ƞ_p_^2^* = .290. However, there were no significant main effects of gloss levels, *F*(2.250, 49.498) = .675, *p* = .530, *ƞ_p_^2^* = .030, or lightness levels, *F*(2.065, 45.428) = .677, *p* = .518, *ƞ_p_^2^* = .030. Additiona l ly, no other significant interactions were found (*p* = .116-.739).

#### Palm Distance

We found that participants moved their hand to a greater extent in the gloss condition (*M_gloss_* = 732.289, *SE* = 84.504) than in the lightness condition (*M_lightness_* = 563.149, *SE* = 65.068, *p* = .012). A repeated-measures 3-way ANOVA revealed a signific ant main effect of the perceptual task, *F*(1,22) = 7.596, *p* = .012, *ƞ_p_^2^* = .257. There were no significant main effects of gloss levels, *F*(3,66) = .432, *p* = .721, *ƞ_p_^2^* = .019, or lightness levels, F(2.011, 44.214) = 1.780, p = .180, *ƞ_p_^2^* = .075. There was a significant interactions between gloss levels and lightness levels, *F*(3.678, 80.916) = 2.671, *p* = .042, *ƞ_p_^2^* = .108. Bonferroni posthoc pairwise comparisons were, however, not significant (Supplementary Figure S2), There were no other significant interactions (*p* = .197-.887).

#### Palm Rotation

Overall, participants rotated the hand more during the gloss judgement (*M_gloss_* = 651.805, *SE* = 66.716) than in the lightness judgement task (*M_lightness_* = 484.463, *SE* = 46.687, *p* = .013). Figure 10 visualised individual palm rotation trajectories for four participants exploring the same stimulus (Gloss/Lightness Level 4) across both tasks. While all spent more time and made larger hand rotations in the gloss condition, their trajectories varied substantially, demonstrating individual differences in exploration strategies even when interacting with the same stimuli. A repeated-measure 3-way ANOVA revealed a significant main effect of the perceptual task, *F*(1,22) = 7.361, *p* = .013, *ƞ_p_^2^* = .251. There were no significant main effects of gloss levels, F(3,66) = 1.042, *p* = .380, *ƞ_p_^2^* = .045, or lightness levels, *F*(2.322, 51.090) = 1.401, *p* = .256, *ƞ_p_^2^* = .060. There was a significant interaction between gloss levels and lightness levels, *F*(9, 198) = 2.383, *p* = .014, *ƞ_p_^2^* = .098, however, Bonferroni posthoc pairwise comparisons were not significant (see Supplementary Figure S3 for details). There were no other significant interactions (*p* = .327-.790).

**Figure 10.**
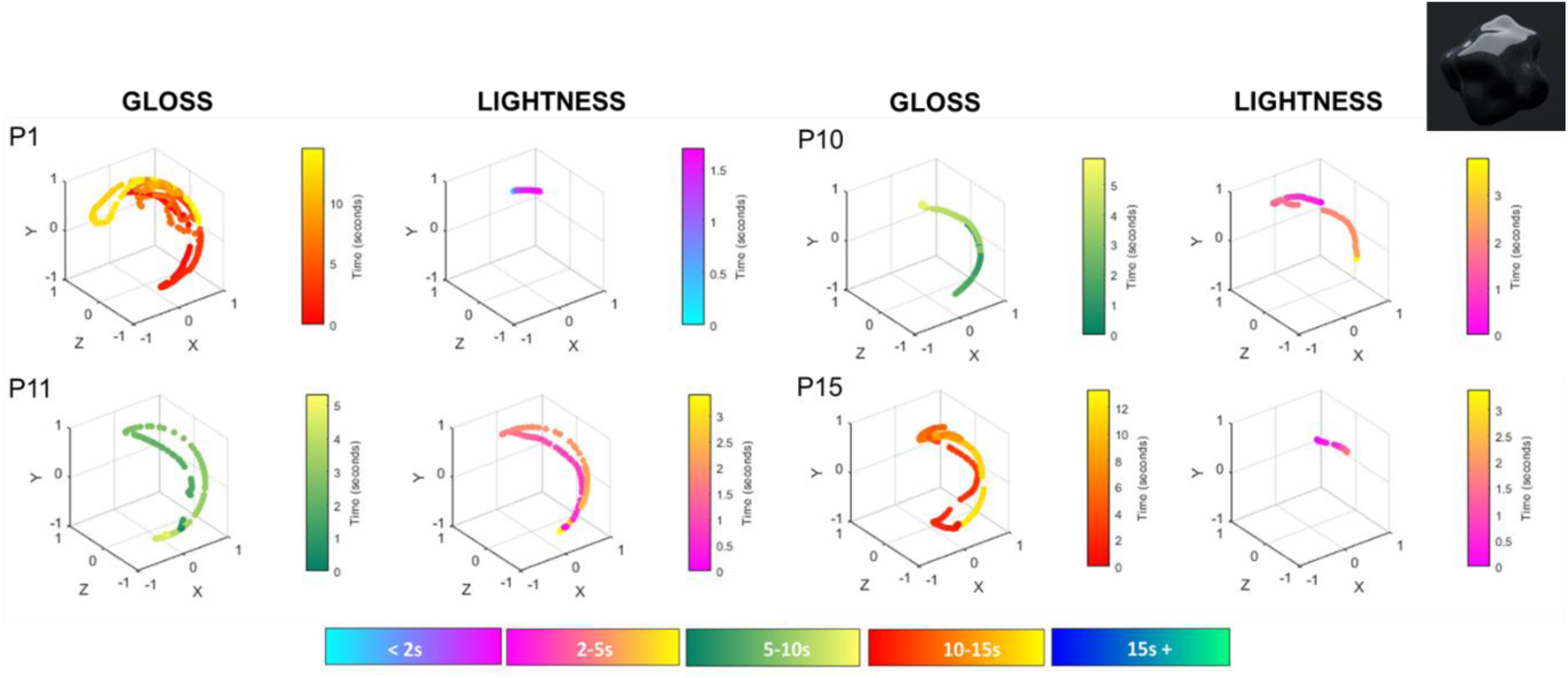
Individual Differences in Palm Rotation Trajectories. The figure shows the palm rotation trajectories of four participants while exploring the same stimulus across both perceptual tasks.

### Observation conditions

#### Looking Time

We did not find a difference in looking time during the gloss judgement task (*M_gloss_* = 6.084, *SE* = .520) and the lightness judgement task (*M*_lightness_ = 5.419, *SE* = .548, *p* = .224). A repeated-measure 3-way ANOVA revealed no significant main effects of perceptual task, *F*(1,22) = 1.563, *p* = .224, *ƞ_p_^2^* = .066, gloss levels, *F*(1.917, 42.174) = .134, *p* = .866, *ƞ_p_^2^* = .006, or lightness levels, *F*(1.901, 41.822) = .348, *p* = .697, *ƞ_p_^2^* = .016. No significant interactions were found (*p* = .501-.873).

### Perceptual Judgements

Overall, stimuli were perceived as slightly glossier when participants interacted with them compared to just observing them. Whereas for lightness judgements, stimuli were perceived as somewhat lighter in the interactive conditions as compared to the observation condition (see Figure 11 for an overview).

**Figure 11.**
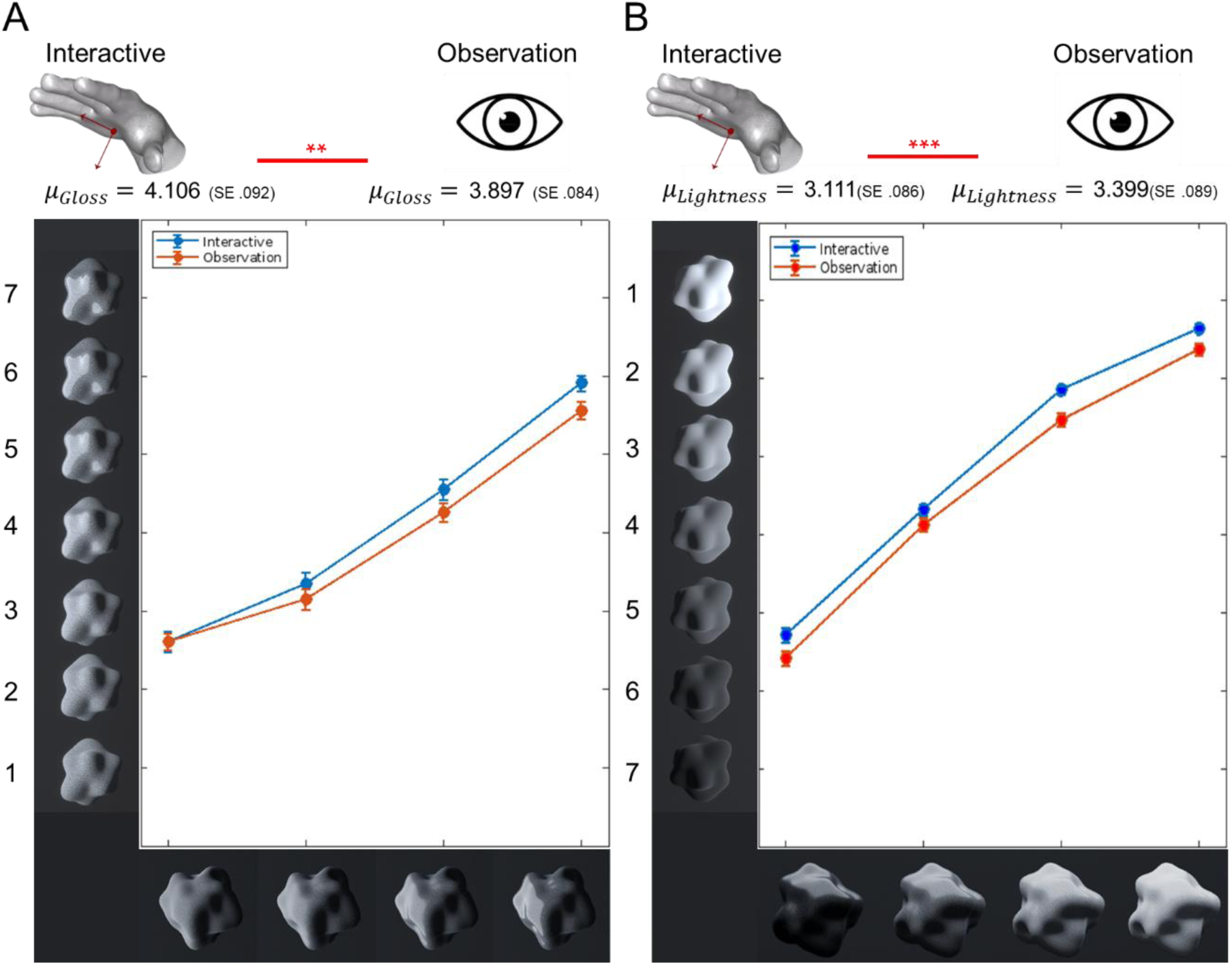
Results perceptual judgements. A) Mean perceived gloss level as a function of stimulus gloss levels, averaged across different lightness levels. B) Mean perceived lightness level as a function of stimulus lightness levels, averaged across different gloss levels. The stimuli on the x-axes are illustrative representations of different gloss/ lightness levels, and are included for visualisation purposes only. The axes of panel B were adjusted so that higher values correspond to lighter levels. Different coloured lines representing the two exploration conditions. Error bars represent 1 ± SE.

#### Gloss perception

A pair-sample t-test showed significant difference in mean perceived gloss, *t*(367) = 2.925, *p* = .004, with stimuli in the interactive condition (*M* = 4.106, *SE* = .092) being rated as more glossy than in the observation condition (*M* = 3.897, *SE* = .084).

#### Lightness perception

A pair-sample t-test demonstrated a significant difference in mean perceived lightness, *t*(367) = −6.077, *p* <.001, with stimuli in the observation condition (*M* = 3.399, *SE* = .089) being rated as darker than in the interactive condition (*M* = 3.111, *SE* = .086).

### Interparticipant correlations and individual differences

We examined agreement among participants in both their perceptual judgments and exploration behaviours. Perceptual ratings were highly consistent across conditions, with correlation coefficients ranging from .677 to .896, whereas correlations for exploration behaviours (e.g., exploration time, palm distance, palm rotation, looking time) were much lower, ranging from −.002 to .077. Furthermore, individual differences in exploration strategies were evident: some participants exhibited strong task-related effects (e.g., more extensive exploration during gloss judgements), while others showed little to no difference across tasks (see Figure 12).

**Figure 12.**
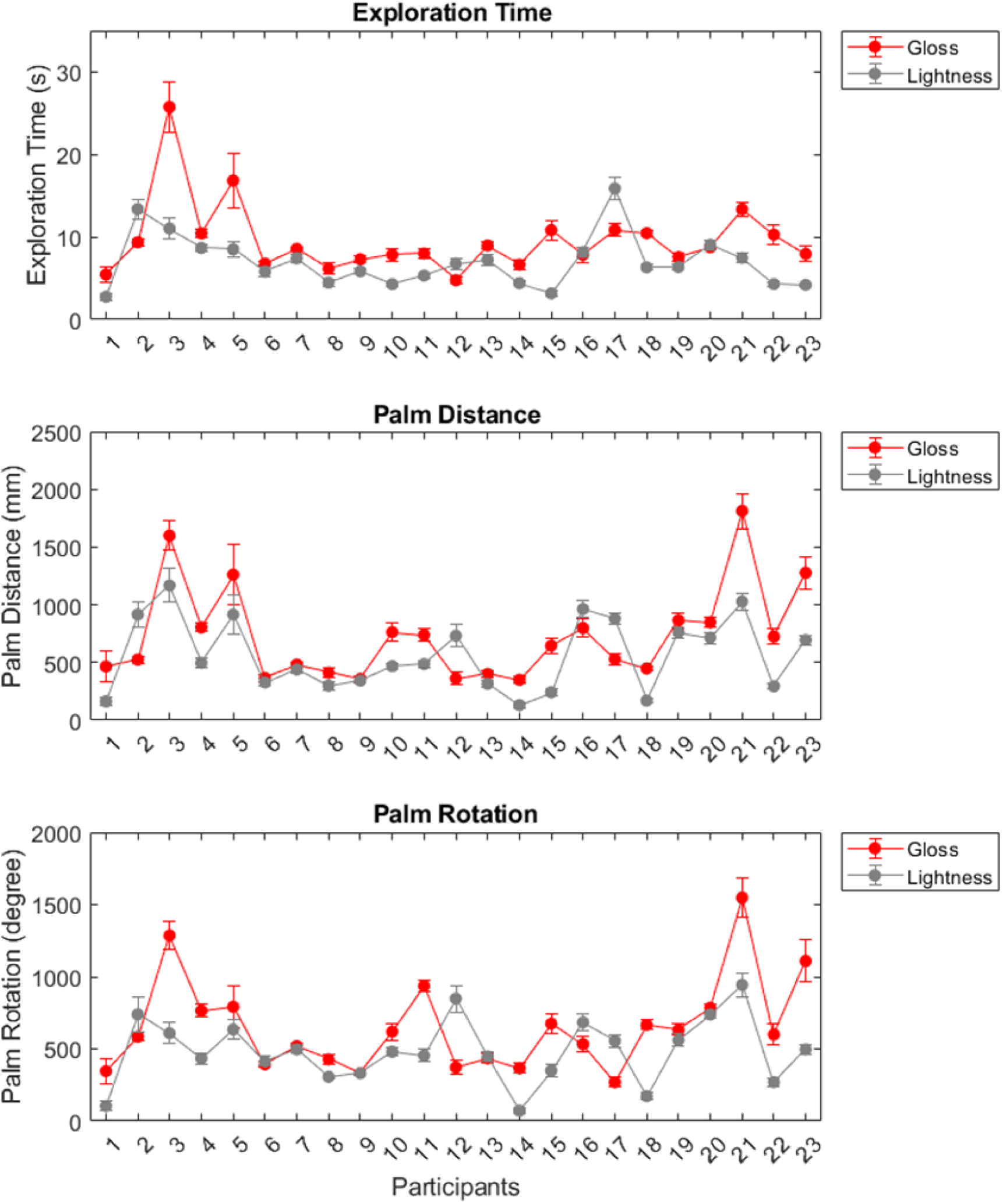
Individual Differences in Exploration strategies across Tasks. A) Mean exploration time, B) Mean palm distance, and C) Mean palm rotation are plotted for each participant. The red line represents exploration during the gloss judgement task, while the grey line represents exploration during the lightness judgement task Error bars indicate ±1 SE. While there is a clear task effect, with greater exploration overall in the gloss condition, participants exhibit highly individualised exploration strategies, as indicated by low interparticipant correlations across tasks. However, when comparing each participant’s exploration behaviour across tasks, correlations were slightly stronger, indicating that in addition to task-dependent differences, participants also showed stable individual exploration tendencies: Exploration time: mean = 0.09 [-0.43,0.43]; Palm Distance: mean = 0.10[-0.30,0.67]; Palm rotation: mean = 0.16 [-0.21,0.58].

## Discussion

The primary aim of this study was to examine whether exploration behaviour during visual material perception is influenced by the specific perceptual task, specifically whether participants adjust their exploration strategies when judging gloss versus lightness. To explore this, we conducted a VR experiment to assess how exploration behaviour is shaped by the task demands (gloss vs. lightness judgments), with participants either interacting with or observing objects before making matching judgments. We assessed observers’ exploration patterns and perceptual judgments. We expected that observers would engage more with the stimulus during gloss judgement tasks since more potential information could be gained by rotating a glossy object, e.g. by generating highlight movement across the objects surface.

### Task-driven exploration behaviour during gloss judgements

We found that participants consistently engaged in more extensive exploration - longer exploration time, greater palm movement, and more rotation, during gloss judgments compared to lightness judgments. These differences in movement patterns could suggest a strategic prioritisation of relevant cues to the perceptual task. For example, for gloss perception, rotation emerges as a key feature of natural object manipulation, reflecting participants’ desire to actively explore objects in a way that generates relevant visual information that facilitates their gloss judgements. In contrast, lightness judgments may rely more on global properties (e.g. brightest area on the object), necessitating less extensive exploration (Toscani et al., 2013 & 2017).

Previous research supports the role of motion in enhancing the perception of surface properties such as gloss. Motion cues provide critical information by allowing dynamic visual features, as such specular highlights, to shift and change as either object or observer moves (Hurlbert, Cumming, & Parker, 1991; Hartung & Kersten, 2002). These visual motion characteristics provided far more about the material properties of the object than static views or serial sampling alone. While humans can perceive gloss from static cues, motion cues enhance gloss perception by improving gloss constancy and allow observers to distinguish between matte and glossy/shiny objects that otherwise appear indistinguishable or ambiguous when static (Wendt et al., 2010; Doerschner et al., 2011; Hartung & Kersten, 2002).

This aligns with our finding that participants engaged in more exploration during gloss judgment tasks, suggesting that observers actively seek out dynamic cues to aid their gloss judgments. However, a question may arise as to the relative contribution of motion and serial sampling in the exploratory behaviours observed in the current study. Serial sampling involves observing a sequence of static views presented in succession, whereas motion introduces temporally coherent dynamic changes that unfold over time (i.e., continuous transformations of object and/or environment). In our study, disentangling these effects would be challenging due to the continuous and interactive nature of virtual reality, which does not allow for frame randomisation to isolate motion effects. Despite this, the broader literature provides evidence for the role of motion in gloss perception. For instance, Sakano and Ando (2010) showed that dynamic cues such as motion parallax enhance perceived gloss. They showed that gloss perception depends not just on motion itself but on the temporal coherence of motion and the corresponding changes in retinal images, which together enhance the perception of gloss. This suggests that disrupting these dynamic cues, such as by scrambling frame order, may impair the visual system’s ability to integrate information about surface properties. While gloss perception can occur from static images, continuous motion appears to play a distinct role in enhancing perceived gloss by maintaining the coherence in the integration of visual cues. In support of this, Bi et al, (2018) demonstrated that scrambling frame order impaired the ability of models trained on motion sequences to predict human perceptual scales for mechanic a l properties. While gloss is a surface property and not a mechanical one, this finding highlights the importance of temporal coherence for interpreting visual information from motion sequences. Our results align with this evidence, as participants’ extensive exploration and rotations during gloss judgments likely reflect an effort to generate motion cues that facilitate gloss perception.

### Exploration behaviour and material appearance

In addition to measuring exploration parameters, we recorded participants’ perceptual judgments of gloss and lightness to ensure that they performed the tasks as intended in the virtual environment. Interestingly, we observed some small yet statistically significant differences in material perception across exploration conditions. While we do not wish to over-interpret these results, we believe it is important to acknowledge and consider them within the broader literature on material perception.

For gloss, stimuli in the interactive conditions were rated slightly glossier than in the observation conditions. Although these effects were weak, they align with existing research suggesting that motion cues, in particular highlight motions – enhance gloss perception (Wendt et al., 2010; Sakano & Ando, 2010). Research has shown that moving objects produce dynamic visual cues such as shifting specular highlights, which are particularly informative for distinguishing glossy from matte surfaces. In which motion-based cues allow the visual system to better detect and interpret surface reflectance properties (Hurlbert, Cumming, & Parker, 1991; Hartung & Kersten, 2002, Doerschner et al. 2011). In our study, the slight increase in perceived gloss during interactive exploration likely reflects participants’ ability to produce and process these dynamic cues through object interaction, thus supporting the importance of motion in gloss perception.

For lightness, objects in the interactive conditions were perceived to be slightly lighter than in the observation conditions. The origin of this effect is less clear, and given its small magnitude, we do not wish to speculate extensively. However, one possible explanation is that the slight variations in motion could have affected the perceived shading or luminance distribution on the object surface, subtly influencing lightness perception. Previous work on lightness constancy suggests that even minor changes in object orientation or observer movement can impact the interpretation of surface shading and luminance gradients, potentially leading to small perceptual shifts in lightness (Toscani et al., 2013; Gilchrist et al., 1999).

While these perceptual differences are small, they suggest that exploration behaviour may subtly modulate material appearance by enhancing certain task-relevant visual cues, particularly in gloss judgments. In which exploratory actions (e.g. rotation and movement) helped observers gather relevant visual information to support their material judgements. Further research should clarify the extent to which interactive exploration meaningfully influences material perception and identify the specific visual cues most impacted by interactive exploration versus mere observation.

### Individual differences

Despite high agreements in their perceptual judgements across conditions (*r* = .677 to .896), participants exhibited considerable individual differences in their exploratory behaviours (*r* = −0.002 to 0.077). This suggests that while participants arrived at similar perceptual conclusions, the underlying exploratory strategies leading to these judgments varied. Participants generally explored more extensively during gloss judgments than lightness judgments, but the extent of this effect varied across individuals. Some participants exhibited strong task-related differences in their exploration, while others showed little distinction between tasks. Figure 12 illustrates this variability, highlighting both task-dependent differences and idiosyncratic exploration strategies.

We did not find a strong stimulus-level driven effect on exploration. While some participants explored glossier objects more than less glossy ones, others showed no difference or even the opposite pattern. This contrasts with the perceptual judgments, such as gloss perception, where nearly all participants judged smoother objects as glossier. As a result, we found high interparticipant correlations for perceptual judgements, but low for exploration parameters. At the same time, however, participants were quite consistent in their overall exploration tendencies. For instance, those who explored more in one perceptual task also tended to explore more in the other, as illustrated by the overall positive intraobserver correlations for exploration parameters across the two perceptual tasks (Figure S4 in Supplementary Materials). Thus, while perception was predominantly modulated by stimulus properties, overall exploration behaviour was mostly driven by the perceptual task. How participants adjusted their explorations within a given task (i.e. across stimulus levels) varied considerably between individuals. These findings support the notion that even though individuals employ different strategies for exploration, they are still able to obtain comparable perceptual information through their individual acts of exploration.

The assumed variability in exploration strategies may stem from attentional biases and task driven exploration, in which individuals prioritise distinct task-relevant features based on their own attentional preferences and strategies. Different participants may, therefore, focus on various surface features, such as specular highlights in gloss judgments or areas of high luminance for lightness judgments, yet arrive at similar perceptual conclusions by extracting information suited to their specific exploration strategies. Research on attentional and visual sampling strategies offers insight into this variability, suggesting that the selection of exploration targets depends on what features participants consider most relevant for the task at hand (Hayhoe & Ballard, 2014). For example, previous studies show that individuals may focus on brighter areas when assessing lightness or prioritise highlight regions during gloss assessments (Toscani, et al., 2013). Despite these differences in exploratory behaviour, individuals could achieve consistent judgments by focusing on the most reliable cues within their preferred areas, supporting convergence in perceptual judgements.

Moreover, given the intrinsic link between attention and gaze (Hoffman & Subramaniam, 1995; Kowler et al., 1995), we know that attentional allocation during visual exploration, which is closely tied to eye movement patterns, varies across individuals. Examining observers’ gaze behaviours under similar paradigms could further reveal how different observers balance attention between regions of interest, yielding similar perceptual information through divergent strategies. Such research could shed light on the individual allocation of attentional resources during active exploration, particularly for visual material judgments.

In sum, participants’ exploratory strategies may differ, yet the goal-driven nature of exploration ensures that they extract the necessary perceptual information to support reliable and consistent judgements. This flexibility in exploration likely enables individuals to adapt to the perceptual demands of the task, resulting in high perceptual agreement despite variability in the underlying exploration patterns.

### Limitation

While VR provides a life-like immersive experimental controllable environment, it is important to acknowledge the limits. E.g. the stimuli used in our experiments, the smooth, computer - generated (CG) objects, may still lack some of the complexity of real-world materials, which often also possess intricate textures, irregular shapes, and diverse optical properties. Artificial stimuli, while effective for isolating certain perceptual variables, may not fully capture the range of cues humans rely on in real-world settings to perceive material properties.

Additionally, exploration behaviour in controlled virtual environments may differ from those in naturalistic settings, where objects interact more dynamically with light and other environmental factors. Moreover, visual material perception is often influenced by haptic feedback, contextual information and texture cues present in naturalistic settings, which could be difficult to simulate accurately in CG environments. The use of simplified stimuli might have limited participants’ ability to engage in the full range of exploratory behaviours that naturalistic stimuli would evoke. Despite these constraints, the use of perlin noise-based textured help introduce naturalistic surface variations, making the stimuli more representative of real-world materials. More importantly, the controlled nature of our study allowed us to systematically isolate the influence of task demands on exploration behaviours, providing insights into the adaptive strategies observers use when judging material properties. Future studies could explore whether these findings extend to more complex, real-world stimuli while maintaining experimental control.

## Conclusion

When given the option, participants rotate objects more during gloss judgements than during lightness judgements, engaging in exploration patterns that enhance the observation of dynamic cues. This supports the role of motion-driven exploratory behaviour in facilitating gloss perception. Our findings, along with existing research, confirm that while humans are capable of perceiving gloss from static visual cues, motion also plays an important role in the perception of visual material properties (Wendt et al., 2010; Doerschner et al., 2011). Our data suggested that individuals utilise a variety of exploratory behaviours to generate information when assessing gloss and lightness, so that different exploration patterns may produce similar perceptual cues crucial for perceiving visual material qualities. Overall, our findings demonstrated the impact of visual material and task in guiding and modulating human interactions with objects.

## Acknowledgement

L.L. and K.D were supported by the Hessisches Ministerium für Wissenschaft und Kunst (HMWK; project ‘The Adaptive Mind’), K.D. and K.D. were supported by the Deutsche Forschungsgemeinschaft (DFG, German Research Foundation) – project number 222641018 – SFB/TRR 135, A5 & B8.

