## supplementary materials for "Exploring For Gloss: Active Exploration in Visual Material Perception"

**Supplementary Figure S1: Average pixel difference analysis of textured bumpy objects with 45-degree rotation**

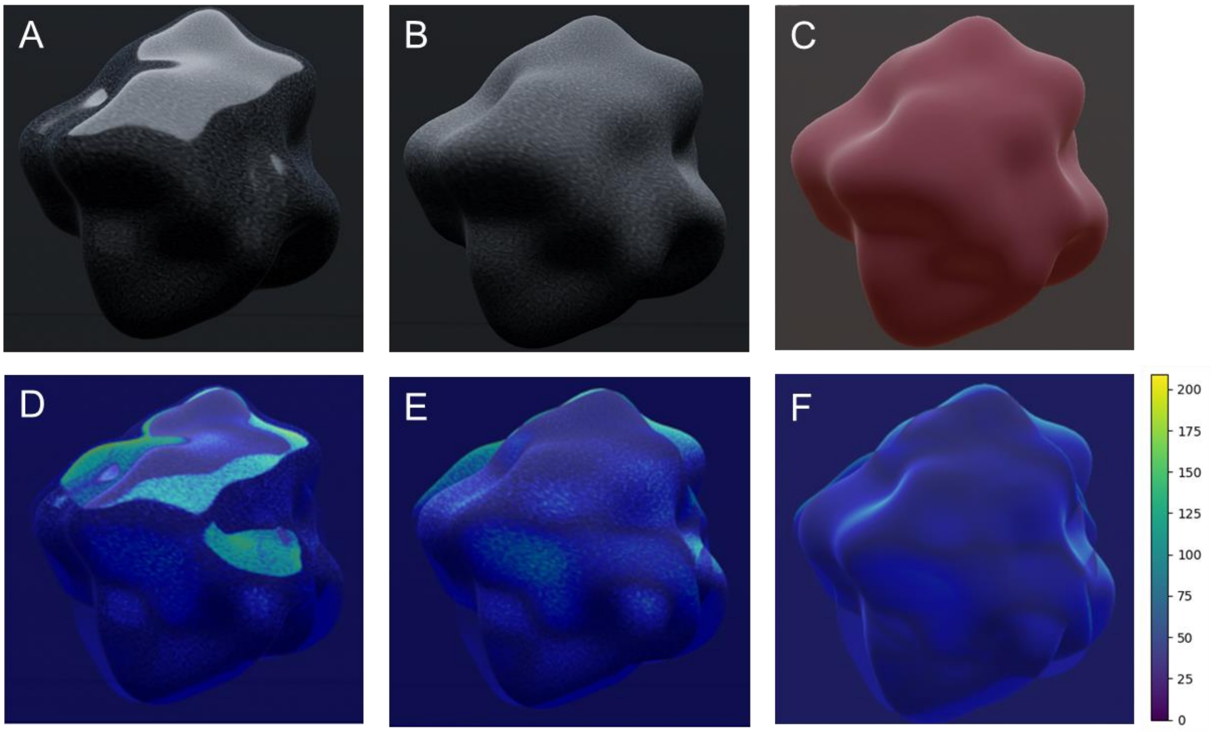


Supplementary Figure S1. **Absolute pixel intensity difference across frames**. Panel A shows a rotating textured high-gloss object, Panel B shows a rotating textured matte object, and Panel C shows a rotating smooth matte object, Panels D, E and F display the average absolute pixel difference across frames, where the objects differ by a 45 degree rotation.

### Supplementary Section A: Pilot experiment - Exploring Differential Exploration Behaviour Across Perceptual Tasks

Before conducting the main experiment, we ran a pilot study to assess whether exploration behaviour differed between perceptual tasks (gloss vs. lightness judgments) and to evaluate the feasibility of our paradigm. The methodology was the same as in the main experiment, except that separate stimulus sets were used for each task.

#### Participants

Twenty-three participants ($M_{age}$ = 26.8, $SD_{age}$ = 3.0, 8 males) from Giessen University participated. All had normal or corrected-to-normal vision and no motor impairments. Three participants were excluded from the analysis due to equipment failure, and one participant was excluded due to non-compliance with the instructions. Participants provided informed consent and were compensated (€8/h).

#### Stimuli and Material

The same 3D shapes from the main experiment were used, but distinct stimulus sets were created for each task.

**Gloss condition:**

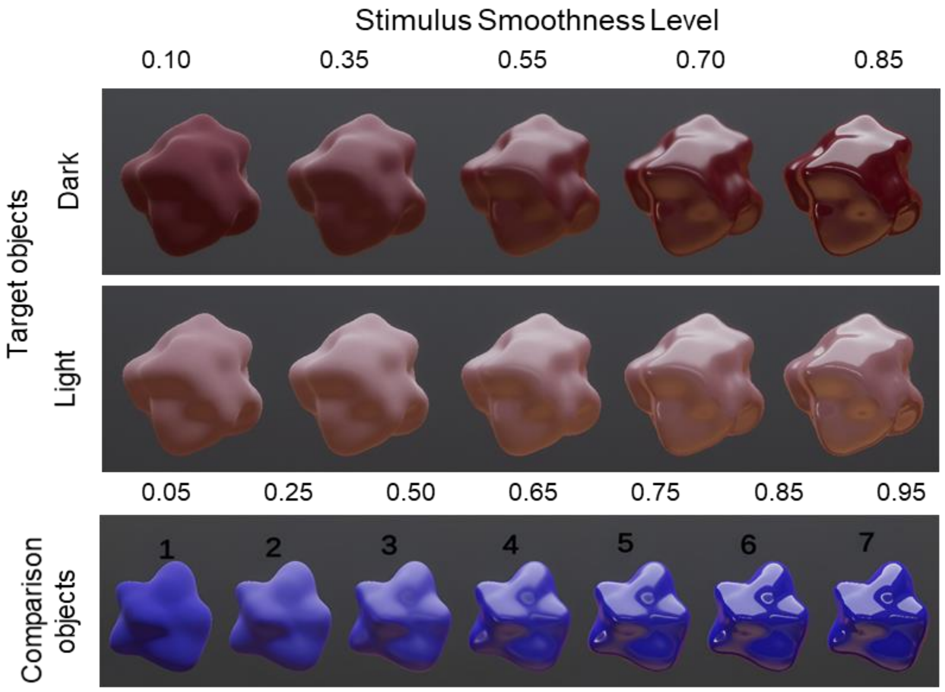


Supplementary Figure A1. Target and comparison objects used in the gloss conditions. Target object: Five gloss levels were generated by varying smoothness values (top panel), with dark and light base colours ([0.208 0.004 0.047] and [0.341 0.243 0.263], respectively). Comparison objects: Seven gloss levels were generated by varying smoothness values (bottom panel).

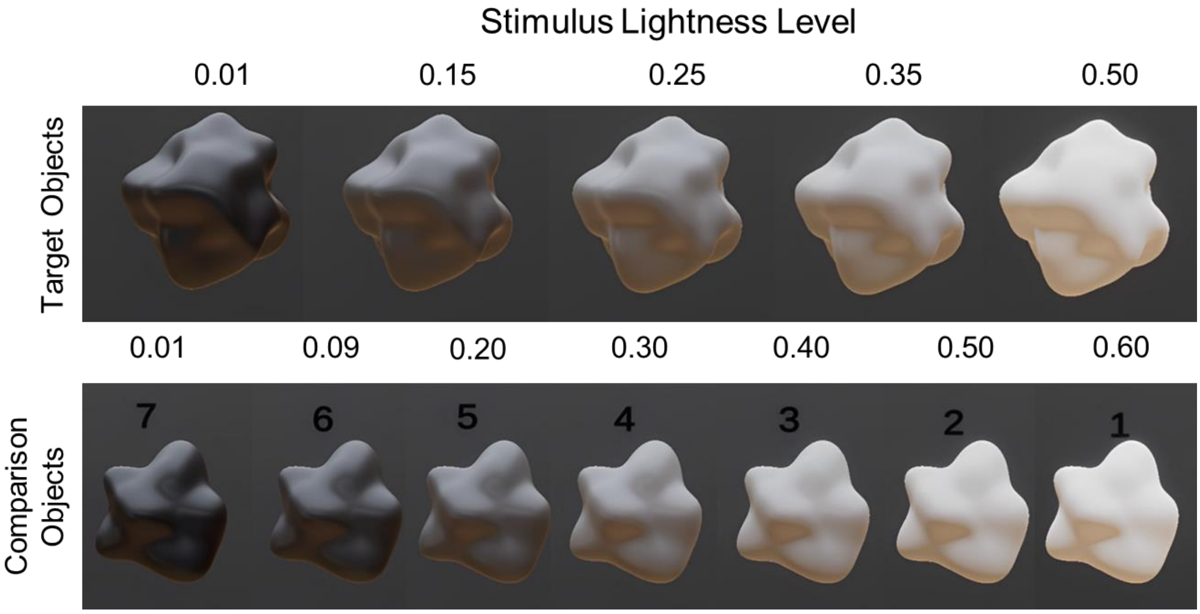


Supplementary Figure A2. Target and comparison objects used in the lightness conditions. Target objects: Five lightness levels were created by adjusting RGB values (top panel). Comparison objects: seven lightness levels were generated by varying RGB values (bottom panel). Smoothness was held constant at 0.5 to prevent strong highlights.

**Lightness condition:**

**Design, Procedure and Measurements**

The virtual environment matched the main experiment, except for a single overhead area light (1721 lumens). In the interactive condition, comparison objects rotated at 1° per frame. The same exploration and hand movement parameters were recorded as in main experiment, including exploration time, looking time, palm distance, and palm rotation.

**Results**

**Perceptual Judgements**

Participants performed the perceptual task sensibly, rating glossier stimuli as glossier and darker stimuli as glossier than lighter ones ($p$ = .003). A small effects of interaction was found, with stimuli appearing slightly glossier in the interactive condition ($p$ = .036). Lightness judgments followed expected patterns, with darker stimuli rated as darker. Participants also perceived stimuli as slightly lighter in the interactive condition ($p$ = .008).

**Exploration Behaviours**

Participants explored longer and made more extensive movements in the gloss condition than in the lightness condition (Supp. Figure A.3).

**Interparticipant correlations and individual differences**

Perceptual judgments (all repetitions considered) were highly consistent across participants ($r$ = .873–.924), while exploration parameters were weakly correlated ($r$ = -.007 to .134), indicating substantial individual differences in how participants explored objects.

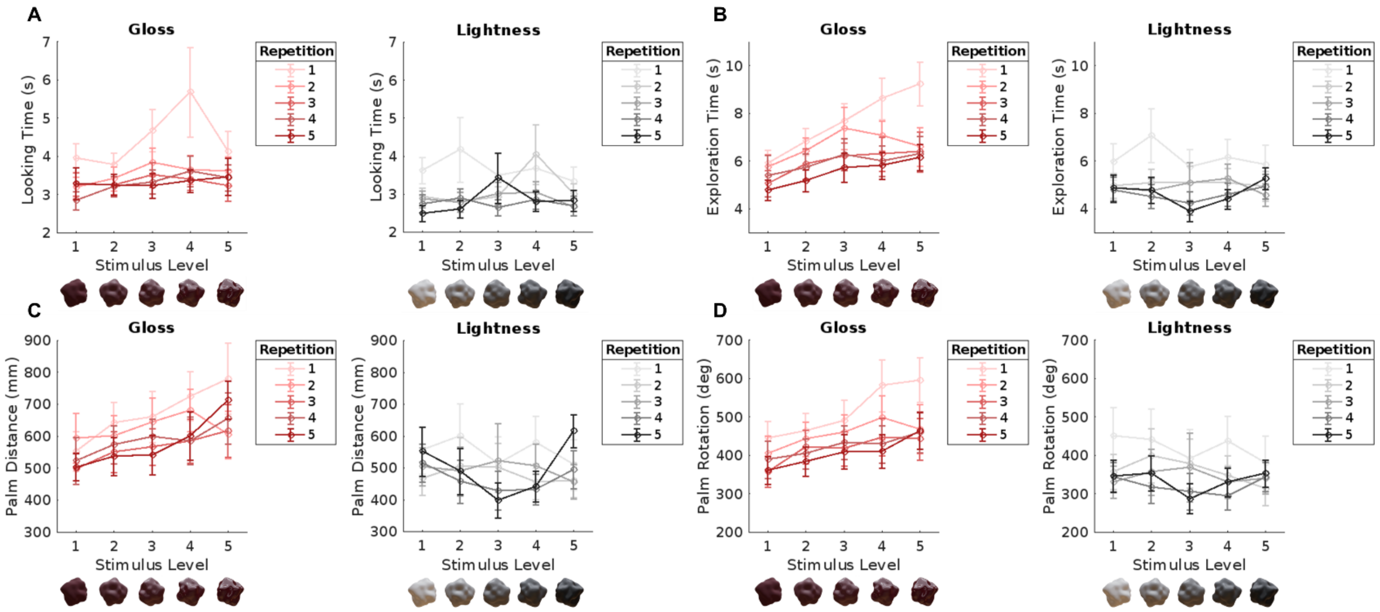


Supplementary Figure A3. **Results exploration parameters.** A) Mean looking time, B) Mean exploration time, C) Mean palm distance, and D) Mean palm rotation across perceptual tasks plotted as a function of stimulus level and repetition. Error bars represent 1 ± SE. Exploration time: significantly longer in gloss (*M* = 6.35) than lightness condition (*M* = 5.08), *p* = .030; Palm Distance: Move movement in gloss (*M* = 605.51) compared to lightness condition (*M* = 498.84), *p* = .039; Palm rotation: More extensive rotation in gloss (*M* = 443.58) than in lightness condition (*M* = 355.98), *p* = .027; Looking time: Longer in gloss (*M* = 3.59) than in lightness condition (*M* = 3.06), *p* = .040.

### Supplementary Tables: Chromaticity and Luminance Measurements of Stimuli Material Used in the Experiment

| Material | Description | x | y | Y |
|---|---|---|---|---|
| Lightness Level 4 + Gloss Level 1 | Darkest + Least Glossy | 0.27229935 | 0.30008465 | 2.6240437 |
| Lightness Level 4 + Gloss Level 2 | | 0.27487105 | 0.30201185 | 2.798888 |
| Lightness Level 4 + Gloss Level 3 | | 0.27151537 | 0.30273432 | 3.2250199 |
| Lightness Level 4 + Gloss Level 4 | | 0.27724054 | 0.30440921 | 2.6911681 |
| Lightness Level 3 + Gloss Level 1 | Medium Dark + Least Glossy | 0.27256659 | 0.30063716 | 5.4658365 |
| Lightness Level 3 + Gloss Level 2 | | 0.27379712 | 0.3016468 | 5.6644392 |
| Lightness Level 3 + Gloss Level 3 | | 0.27439976 | 0.30233642 | 5.6458001 |
| Lightness Level 3 + Gloss Level 4 | | 0.27447861 | 0.30221501 | 5.8371167 |
| Lightness Level 2 + Gloss Level 1 | Medium Light + Least Glossy | 0.27491757 | 0.3027266 | 9.784565 |
| Lightness Level 2 + Gloss Level 2 | | 0.27407318 | 0.30186909 | 9.6434231 |
| Lightness Level 2 + Gloss Level 3 | | 0.27400523 | 0.30181307 | 9.6919327 |
| Lightness Level 2 + Gloss Level 4 | | 0.27364811 | 0.30144465 | 9.386096 |
| Lightness Level 1 + Gloss Level 1 | Lightest + Least Glossy | 0.27792901 | 0.30468288 | 14.814525 |
| Lightness Level 1 + Gloss Level 2 | | 0.27782443 | 0.30455044 | 14.784842 |
| Lightness Level 1 + Gloss Level 3 | | 0.27748197 | 0.30425906 | 14.415988 |
| Lightness Level 1 + Gloss Level 4 | | 0.27737692 | 0.30405116 | 14.289672 |
| Gloss Comparison Level 1 | Least Glossy | 0.27244398 | 0.30094627 | 5.8149095 |
| Gloss Comparison Level 2 | | 0.27135217 | 0.29983509 | 6.2336197 |
| Gloss Comparison Level 3 | | 0.27441928 | 0.3016746 | 7.7926064 |
| Gloss Comparison Level 4 | | 0.27381656 | 0.30175465 | 6.5047703 |
| Gloss Comparison Level 5 | | 0.27441522 | 0.30243209 | 6.691278 |
| Gloss Comparison Level 6 | | 0.2744728 | 0.30233312 | 6.682951 |
| Gloss Comparison Level 7 | Most Glossy | 0.27382568 | 0.30131453 | 7.6121249 |
| Lightness Comparison Level 1 | Lightest | 0.27731583 | 0.30436316 | 16.094997 |
| Lightness Comparison Level 2 | | 0.27553201 | 0.30244714 | 10.584188 |
| Lightness Comparison Level 3 | | 0.27043933 | 0.29886231 | 6.4705005 |
| Lightness Comparison Level 4 | | 0.27183971 | 0.30044967 | 4.5398126 |
| Lightness Comparison Level 5 | | 0.27098373 | 0.30022386 | 2.3966544 |
| Lightness Comparison Level 6 | | 0.27669021 | 0.30364037 | 0.71538365 |
| Lightness Comparison Level 7 | Darkest | 0.27930307 | 0.31002578 | 0.36365765 |

Supplementary Table S1. **Chromaticity and Luminance Measurements of Stimuli Material Using Target Object Mesh.** Measured chromaticity and luminance values for each material used in the experiment are presented. These measurements were conducted on a 3D object mesh identical to those employed for the target object. *Note*: Object position: Vector3 (56.485, 3.498, 1.660); Camera position: Vector3 (56.450, 2.406, 1.569).

| Material | Description | x | y | Y |
|---|---|---|---|---|
| Lightness Level 4 + Gloss Level 1 | Darkest + Least Glossy | 0.27133119 | 0.29890999 | 3.3763936 |
| Lightness Level 4 + Gloss Level 2 | | 0.27259284 | 0.3000378 | 3.4732652 |
| Lightness Level 4 + Gloss Level 3 | | 0.27423647 | 0.3016592 | 3.2702694 |
| Lightness Level 4 + Gloss Level 4 | | 0.27734661 | 0.30366871 | 2.3683012 |
| Lightness Level 3 + Gloss Level 1 | Medium Dark + Least Glossy | 0.2710349 | 0.29887974 | 3.493552 |
| Lightness Level 3 + Gloss Level 2 | | 0.2716724 | 0.29932061 | 4.1918888 |
| Lightness Level 3 + Gloss Level 3 | | 0.27212492 | 0.29972175 | 3.717963 |
| Lightness Level 3 + Gloss Level 4 | | 0.27272111 | 0.29994819 | 3.3258269 |
| Lightness Level 2 + Gloss Level 1 | Medium Light + Least Glossy | 0.26985517 | 0.29663208 | 7.4018989 |
| Lightness Level 2 + Gloss Level 2 | | 0.26735586 | 0.29901186 | 5.1115179 |
| Lightness Level 2 + Gloss Level 3 | | 0.27068886 | 0.29768378 | 7.0473428 |
| Lightness Level 2 + Gloss Level 4 | | 0.27192929 | 0.29864329 | 6.2265134 |
| Lightness Level 1 + Gloss Level 1 | Lightest + Least Glossy | 0.27573466 | 0.30201897 | 12.949505 |
| Lightness Level 1 + Gloss Level 2 | | 0.27602714 | 0.3024202 | 13.510004 |
| Lightness Level 1 + Gloss Level 3 | | 0.27614042 | 0.30259603 | 13.144261 |
| Lightness Level 1 + Gloss Level 4 | | 0.27588165 | 0.30202335 | 12.401702 |
| Gloss Comparison Level 1 | Least Glossy | 0.2720688 | 0.29924855 | 5.5104618 |
| Gloss Comparison Level 2 | | 0.27087697 | 0.29803878 | 3.8712254 |
| Gloss Comparison Level 3 | | 0.27101254 | 0.29881516 | 3.751112 |
| Gloss Comparison Level 4 | | 0.27151346 | 0.30023345 | 3.1446123 |
| Gloss Comparison Level 5 | | 0.2720938 | 0.29941979 | 4.609446 |
| Gloss Comparison Level 6 | | 0.27220976 | 0.29918823 | 3.1390398 |
| Gloss Comparison Level 7 | Most Glossy | 0.27251744 | 0.29942572 | 3.196631 |
| Lightness Comparison Level 1 | Lightest | 0.27484778 | 0.29861164 | 15.454765 |
| Lightness Comparison Level 2 | | 0.27380574 | 0.30393234 | 9.3071146 |
| Lightness Comparison Level 3 | | 0.27064842 | 0.30041999 | 4.4755797 |
| Lightness Comparison Level 4 | | 0.26882419 | 0.29629856 | 3.8305645 |
| Lightness Comparison Level 5 | | 0.26653776 | 0.29682744 | 4.3711867 |
| Lightness Comparison Level 6 | | 0.27040097 | 0.29974431 | 1.1084398 |
| Lightness Comparison Level 7 | Darkest | 0.28123486 | 0.30627042 | 0.53237265 |

Supplementary Table S2. **Chromaticity and Luminance Measurements of Stimuli Material Using a Flat Cube Mesh.** Measured chromaticity and luminance values for each material used in the experiment are presented. These measurements were conducted on a flat cube mesh. *Note*: Object position: Vector3 (56.489, 3.476, 1.784); Camera position: Vector3 (56.491, 2.314, 1.741).

### Supplementary Figure S2-3: interaction between gloss and lightness levels for palm

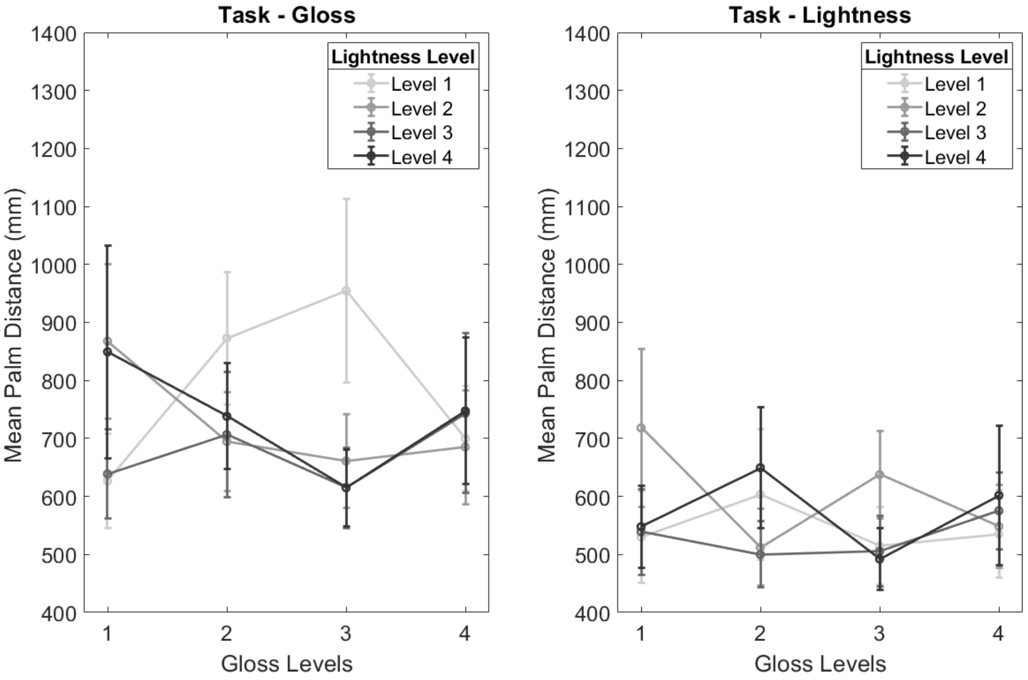


Supplementary Figure S2 **Interaction between gloss and lightness levels on palm distance.** Left panel) Mean palm distance during the gloss judgement task, plotted as a function of gloss level and lightness level. Right panel) Mean palm distance during the lightness judgement task, plotted as a function of gloss level and lightness level. Error bars represent 1 ± SE. As illustrated in the figure, while there was a significant interaction between gloss and lightness levels, they do not alter the overall conclusion of the analysis.

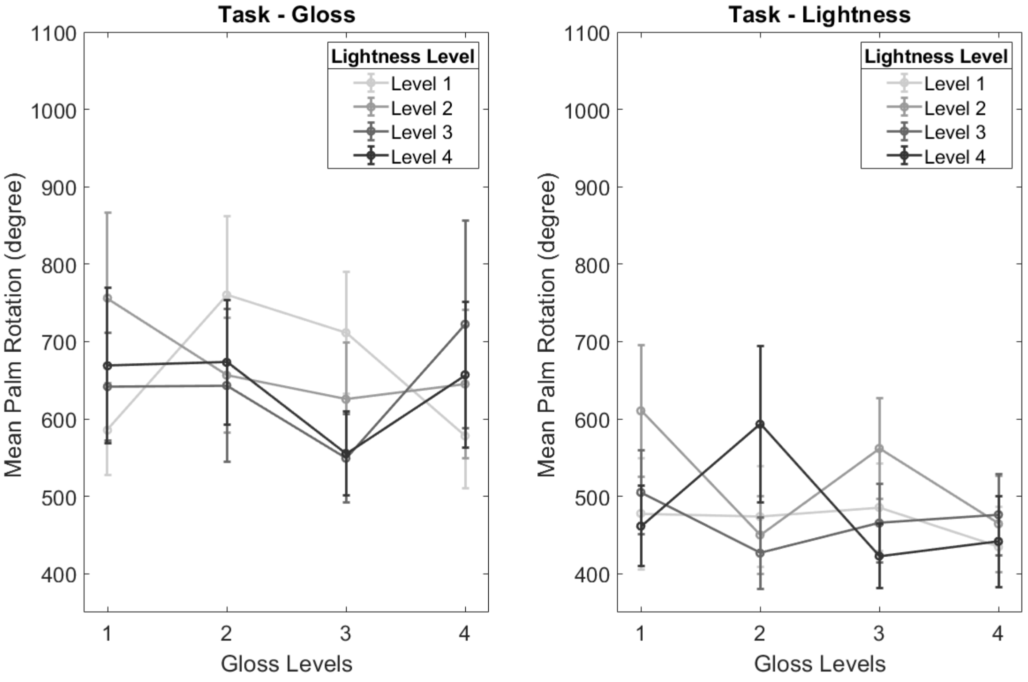


Supplementary Figure S3. **Interaction between gloss and lightness levels on palm rotation.** Left panel) Mean palm rotation during the gloss judgement task, plotted as a function of gloss level and lightness level. Right panel) Mean palm rotation during the lightness judgement task, plotted as a function of gloss level and lightness level. Error bars represent 1 ± SE. As illustrated in the figure, although a significant interaction between gloss and lightness levels was observed, the post-hoc contrasts were not significant.

### Supplementary Figure S4: Individual consistencies in exploration across tasks

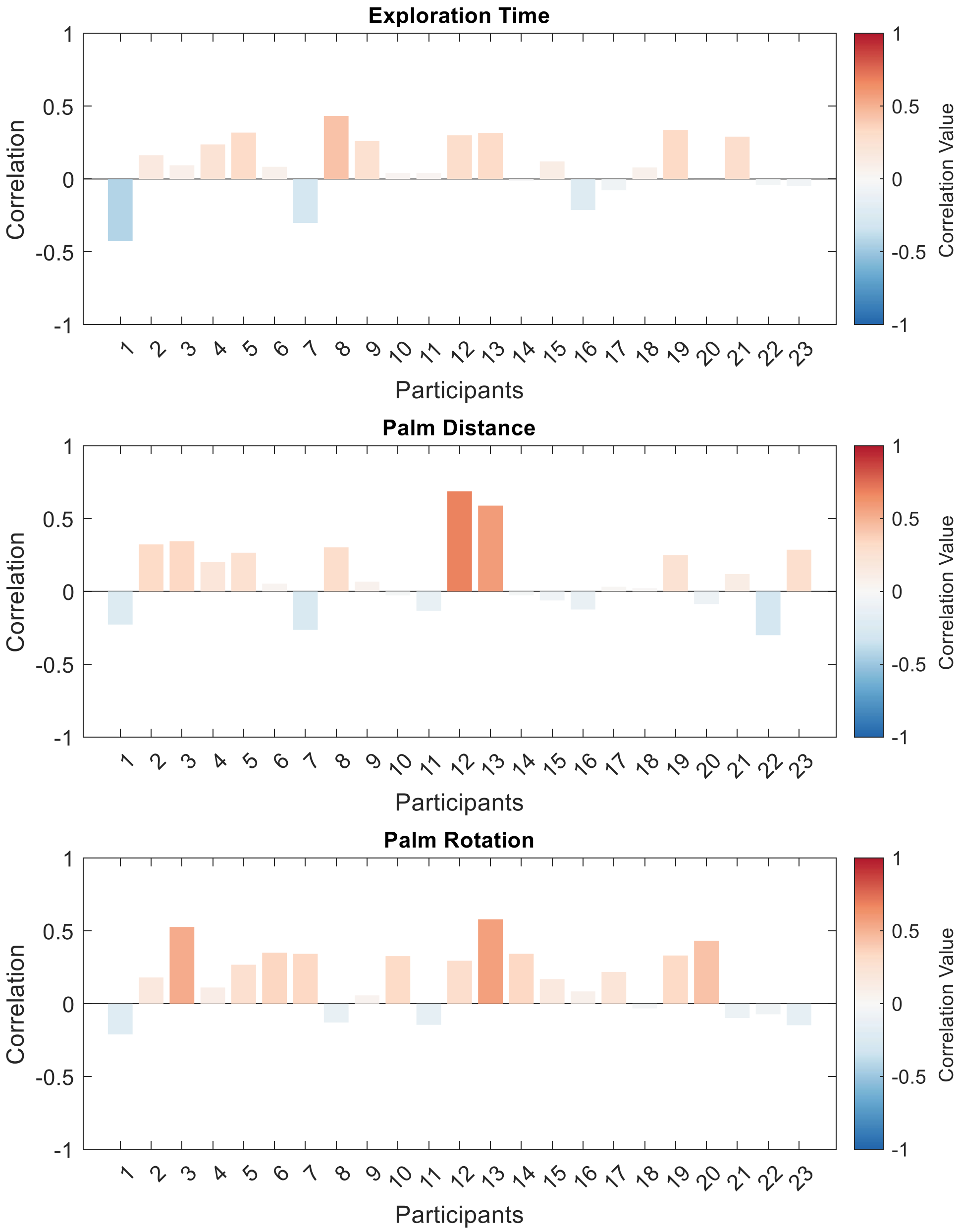


Supplementary Figure S4. **Correlation of Exploration Parameters Across Tasks.** This figure presents the correlation between gloss and lightness exploration parameters for each participant, illustrating individual consistency in exploration strategies across tasks. Each bar represents the correlation for a single participant, with warmer colours indicating stronger positive correlations and cooler colours representing negative correlations Several, but not all, participants exhibited stable exploration tendencies across tasks.
